# Resilience in a time of stress: revealing the molecular underpinnings of coral survival following thermal bleaching events

**DOI:** 10.1101/2024.04.02.587798

**Authors:** Brook L. Nunn, Tanya Brown, Emma Timmins-Schiffman, Miranda Mudge, Michael Riffle, Jeremy B. Axworthy, Jenna Dilwort, Carly Kenkel, Jesse Zaneveld, Lisa J. Rodrigues, Jacqueline L. Padilla-Gamiño

## Abstract

Coral bleaching events from thermal stress are increasing globally in duration, frequency, and intensity. Bleaching occurs when a coral’s algal symbionts are expelled, resulting in a loss of color. Some coral colonies survive bleaching, reacquire their symbionts and recover. In this study, we experimentally bleached *Montipora capitata* colonies to examine molecular and physiological signatures of intrinsic differences between corals that recover (resilient) compared to those that die (susceptible). All colonies were collected from the same bay and monitored for eight months post-bleaching to identify specific colonies exhibiting long-term resilience and survival. Using an integrated systems-biology approach that included quantitative mass spectrometry-based proteomics, 16S rRNA of the microbiome, total lipids, symbiont density and diversity, we explored molecular-level mechanisms of tolerance in pre- and post-bleached colonies and found biomarkers of resilience that can confidently identify resilient and susceptible corals before thermal-induced bleaching events. Prior to thermal stress, resilient corals were characterized by a more diverse microbiome and increased abundances of proteins involved in multiple carbon and nitrogen acquisition strategies, symbiont retention and acquisition, and pathogen resistance. Susceptible corals had early signs of symbiont rejection and had resorted to utilizing urea uptake pathways for carbon and nitrogen. Further, molecular signatures identified prior to bleaching were amplified after bleaching, suggesting these pathways may be deterministic in a colony’s fate. Our results have important implications for the future of reefs, revealing molecular factors necessary for survival through thermally-induced bleaching events and providing diagnostic biomarkers for coral reef management.

**Significance statement:** Corals are being negatively impacted by the increase in the number and duration of thermal-induced bleaching events. There are, however, some individuals within a single species that will bleach and, after time, reacquire symbionts and physiologically recover while neighboring colonies will die. Here, we used a multidisciplinary approach to understand the biochemical details of the physiological changes of resilient and susceptible *Montipora capitata* to thermal-induced bleaching. Resilient corals were characterized by their use of multiple carbon and nitrogen acquisition strategies, metabolically active symbiont relationships, abundant antiviral proteins, and a diverse microbiome. We reveal a multi-factor molecular-level approach for confidently identifying resilient and susceptible coral colonies so that environmental managers can rapidly select quality candidates for propagation while in the field.

## Introduction

Coral reefs are one of the most diverse and structurally complex ecosystems on Earth, providing shelter and habitat for many organisms (1). Tens of millions of people in more than one hundred countries have coastlines adjacent to coral reefs and depend on them for their livelihoods (2). Unfortunately, coral reefs are declining rapidly throughout the world due to pollution, coastal development, overexploitation (3, 4), and effects associated with climate change (5, 6). As global seawater surface temperatures increase, large-scale thermal-induced coral bleaching events (loss of distinctive coral coloration due to expelling of algal symbionts) are becoming increasingly common worldwide (7–9).

When water temperatures surpass a thermal threshold for a given coral species it leads to a breakdown in the association between the coral host and the symbiotic algae, Symbiodiniaceae. This breakdown results in symbiont expulsion and is termed “coral bleaching” due to the loss of the pigmented symbiont (3). In addition to the distinct colors, Symbiodiniaceae provide much of the energetic requirements for the host in the form of organic carbon and nitrogen photosynthetic byproducts (10). The expulsion of Symbiodiniaceae during bleaching events metabolically compromises the host (11), leading to reduced physiological performance, reduced reproductive capacity and can lead to widespread mortality.

A coral colony is a holobiont – a collection of organisms that includes the coral, its symbionts, and the microbial microbiome that exists on and in coral tissue. The diversity of the microbiome on corals frequently decreases with thermal-induced bleaching (12, 13). While some of these changes may be associated with the loss of Symbiodiniaceae, other changes in microbiome diversity appear to be driven by thermal stress directly (14). If cascading microbiome changes induced by bleaching disrupt relationships with bacteria or archaea that benefit coral host metabolism or pathogen defense, then those microbiome changes may also contribute to post-bleaching coral mortality.

Coral microbiomes demonstrate host specificity, bolstering the hypothesis that bleaching-induced microbiome disruptions play a significant role in host mortality (15–19). However, disentangling relationships between corals and their microbiomes is challenging without gnotobiotic models. Moreover, although genomic evidence and correlations between taxonomic abundance and disease provide some hints, it is still unclear which specific microbiome changes associated with bleaching or other stressors are helpful or harmful for the survival of the coral host. Indeed, while a rich literature documents alterations to microbiome structure and stability by many specific stressors and coral diseases — including changes to beta-diversity (20, 21), richness (22), and the abundance of particular taxa — far fewer data are available on which of these diverse microbiome changes best predict the future survival of coral hosts. Thus, data linking microbiome change with subsequent coral survival could be vital to interpreting the ecological consequences of shifts in bleaching-induced microbiome structure for corals.

Coral death due to the interconnected physiological and microbiological consequences of bleaching events can lead to total community collapse if mortality is widespread. However, some coral species, and individuals within species, are resilient to the effects of bleaching and appear to return to pre-bleaching status. Corals resilient to thermal stress can reacquire symbionts, fully recover physiological performance and yield viable gametes (e.g., 23), though the timescale for post-bleaching recovery can vary from weeks to a year (24–27). Post-bleaching recovery times may be shorter in corals that have previously been exposed to multiple annual bleaching events (28), perhaps by priming the physiological responses necessary to survive bleaching.

There are several possible contributing factors that may provide greater coral holobiont resilience to bleaching events: host- and/or symbiont-species (29–32), host genotype (33–38), or the microbial constituents of the holobiont’s microbiome (39, 40). At the center of this complex equation is the host’s metabolic capacity before and after bleaching (i.e., how is the coral acquiring nutrients to sustain growth and immune function?). Without adequate carbon and nitrogen, the coral cannot maintain systems of cellular and tissue repair, retain, or reacquire symbionts, or fight off pathogens (e.g.,41, 42). This suggests that the pre-bleaching molecular physiology of the coral host may be the most important factor in determining resilience to bleaching stress and mortality.

The interactions between an organism and its environment are complex and depend upon ecological and evolutionary history. Corals in Hawai’i, Kāneo’he Bay, O’ahu have endured an increased frequency of bleaching events. Results from bleaching surveys in the bay have shown a decrease in the proportion of coral colonies bleached over time (62% in 1996 to 30% in 2015) but an increase in the number of colonies dying (<1% in 1996 to 22% in 2015). Kāneo’he Bay is inhabited by the reef-building Scleractinian coral *Montipora capitata* (43). These corals are typically found in tropical waters, living within 1-2°C of their upper thermal limit (44, 45). This suggests that projected future water temperatures will specifically threaten this species (e.g., 46). *Montipora capitata* has been shown to have higher thermal tolerance than other Hawaiian coral species (47) and is thereby an ideal candidate to study the long-term effects of thermal acclimatization or adaptation to reveal the underlying molecular physiology that supports bleaching resilience.

In this study, we report the results of a joint coral physiology, proteomic, lipid, and microbiome analysis comparing the features of *M. capitata* coral colonies that recovered *in situ* from experimental bleaching against the features of those that died. Colonies of visually healthy *M. capitata* were exposed to 30°C water in experimental tanks for three weeks to simulate thermal stress and induce bleaching. Samples for proteomics, microbiome diversity, total lipids, Symbiodiniaceae density and clade diversity were collected before thermal stress (T_1_) and three weeks after bleaching occurred (T_2_) (Fig.1). A control set of the same colonies did not undergo thermal-induced bleaching. After the three weeks, corals were outplanted to the field and monitored for eight months to identify colonies that recovered from the thermal-induced bleaching event and those that died. These outcomes were then retroactively used to label the previously collected samples as deriving from *resilient* or *susceptible* colonies. The proteomic, microbial, and physiological differences uncovered in this study thus describe intraspecific differences associated with bleaching resilience in the field. Because coral colonies were collected from and outplanted to the same location at Moku O Lo’e island in Kāneo’he Bay, they experienced the same water and thermal conditions throughout the study.

**Fig. 1.**
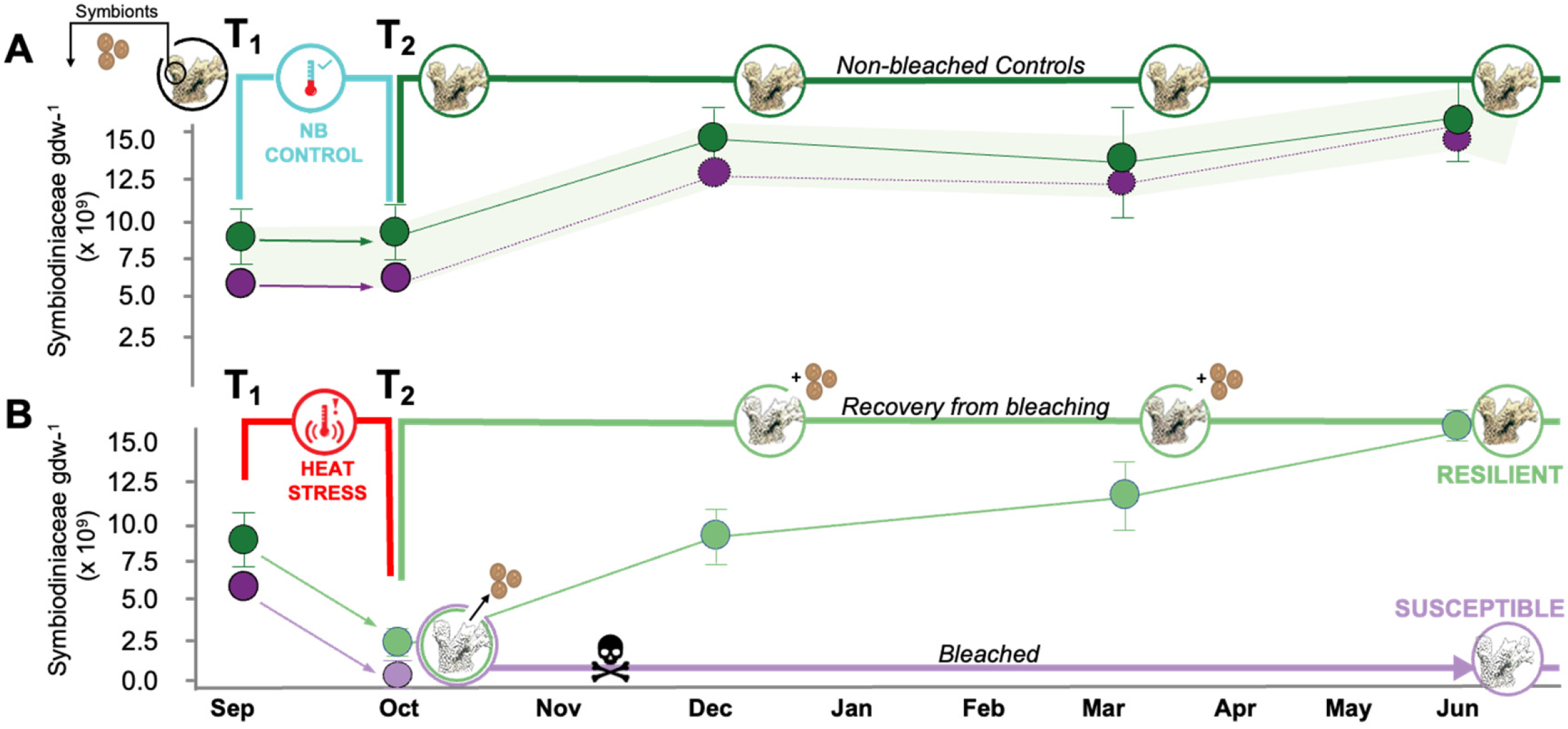
**A:** Total cell counts (x10^9^) for Symbiodiniaceae per gdw^-1^ (*n=*6 for each point) illustrating the experimental design to track effects of thermally induced bleaching on 12 *Montipora capitata* colonies that were monitored for 8 months. After 8 months, each colony was retroactively labeled and 6 colonies that reacquired symbionts and recovered (green) and 6 colonies that were susceptible to thermal stress (purple) were characterized. Symbiodiniaceae density for the (**A**) control cohort maintained at ambient (25°C) temperature and (**B**) experimental cohort that underwent thermally induced bleaching (30°C) for 4 weeks; light green: (*n=*6) resilient colonies reacquired symbionts and recovered post-bleaching; light purple: (*n=*6) colonies susceptible to thermally induced bleaching that did not recover. Coral sub-samples were collected before (T_1_) and after exposure to thermal stress (T_2_) to assess host performance and symbiont and microbial composition in corals. Note that resilient and susceptible colonies were identified three months later (December); after this period no additional mortality was observed.

Comparison of resilient vs. susceptible *M. capitata* using semi-quantitative proteomics allowed the generation of metabolic maps of abundant enzymes and outlined coral’s energetic priorities that may confer resilience in a changing climate. Several proteins significantly differed between resilient and susceptible colonies prior to experimental thermal stress, contributing to bleaching resilience. These resilience-associated proteins reveal dominant nutritional and metabolic strategies underpinning the ability to survive bleaching. The proteome also revealed evidence for symbiont rejection, antiviral activity, enhanced immune response to pathogens, and carbon and nitrogen pathways exclusive to resilient colonies. Additional molecular-level metrics of the holobionts were monitored, including total lipids, microbiome diversity, and symbiont density and diversity, allowing us to evaluate and establish whole-organism biomarkers of resilience. These molecular-level signatures could be used to predict coral resilience or susceptibility prior to a bleaching event. Last, we report a multi-factor approach to identify the corals that will survive future bleaching events, the linchpin to coral management, propagation efforts, and restoration success.

## Results

In September 2017 seventy-four colonies of *Montipora capitata* were tagged (with an ID) and collected, acclimated in tanks for two weeks, sampled (T_1_), gradually exposed to increasing water temperatures to reach 30°C, and held at 30°C for three weeks to simulate a thermal bleaching event (Fig. 1). Water temperatures were returned to ambient temperatures over 4 days, after 24 hrs at that temperature corals were sampled (T_2_) and then were monitored for long-term survival and recovery for eight months (Figs. 1 and S1). After three months (December), 22 colonies died; these colonies will be referred to as “susceptible”. Fifty-two colonies recovered and reacquired symbionts; these colonies will be referred to as “resilient”. After December, no coral mortality was observed. By May, all the colonies that survived reached pre-bleaching coloration (Fig. 1B). At T_1_ and T_2_, we obtained coral samples to examine physiological performance and recovery. Six of the 52 resilient colonies and six of the 22 susceptible colonies were randomly selected for the study and frozen sub-samples at both timepoints were used for mass spectrometry-based proteomics, total lipid content, Symbiodiniaceae density and diversity, and bacterial community composition.

At T_1_, no significant difference was measured in symbiont density between the resilient and susceptible colonies (Fig. 1B, Dataset S1A). During the thermal event, the symbiont density decreased in both cohorts at the same rate; within 3 months after T_2_, the resilient cohort returned to pre-bleaching Symbiodiniaceae density. Additionally, there was no significant impact of bleaching, time point, bleaching tolerance, or colony of origin on symbiont clade abundances (Fig. S1B; Dataset S1B).

A total of 2,193 coral proteins were identified at T_1_, 2,161 coral proteins were identified at T_2_, and 1,424 coral proteins were shared between those timepoints, indicating constitutive expression (Fig. S2, Dataset S2A-G). Analysis of the resilient and susceptible colonies independent of timepoint revealed 2,276 proteins detected in the resilient cohort and 2,066 proteins in susceptible colonies.

### Biological enrichment analysis

Enrichment analysis of Gene Ontology (GO) terms was completed using MetaGOmics (48), an unbiased method to identify significantly different metabolic processes represented in proteomes of the resilient vs. susceptible cohorts at both timepoints using log-fold change (base 2; LFC) of GO term assignments (Dataset S3A-B). Resilient colonies at T_1_ were characterized by multiple cellular responses to external signals, receptor activity, and monosaccharide binding. Sterol esterase activity was the most enriched term (LFC=5.0; Fig. 2A). Six GO terms were significantly enriched in susceptible colonies prior to bleaching (T_1_S) and included proteins involved in urea and amide catabolism, nickel binding, and the removal of superoxides (Fig. 2A). After thermal bleaching, 51 GO terms were enriched in resilient corals, 33 of which were unique (Fig. 2B). Proteomes of resilient colonies post-bleaching (T_2_R) were enriched in regulation of phagocytosis and meiotic cell cycle, vesicle-mediated transport, hormone regulation, and cardiac muscle processes. Although some of the labels for these processes may not seem to apply to corals, proteins associated with the “cardiac muscle process” GO term are involved in sodium and calcium exchange while “regulation of systemic arterial blood pressure” proteins include sodium-driven chloride bicarbonate exchange proteins involved in pH regulation. Many of the identified GO terms in susceptible colonies post-bleaching involved the cellular processing of metabolites, including sterols, methionine, betaine, sarcosine, lipids, and glutamate. Additionally, several terms were related to DNA or RNA processing.

**Fig. 2.**
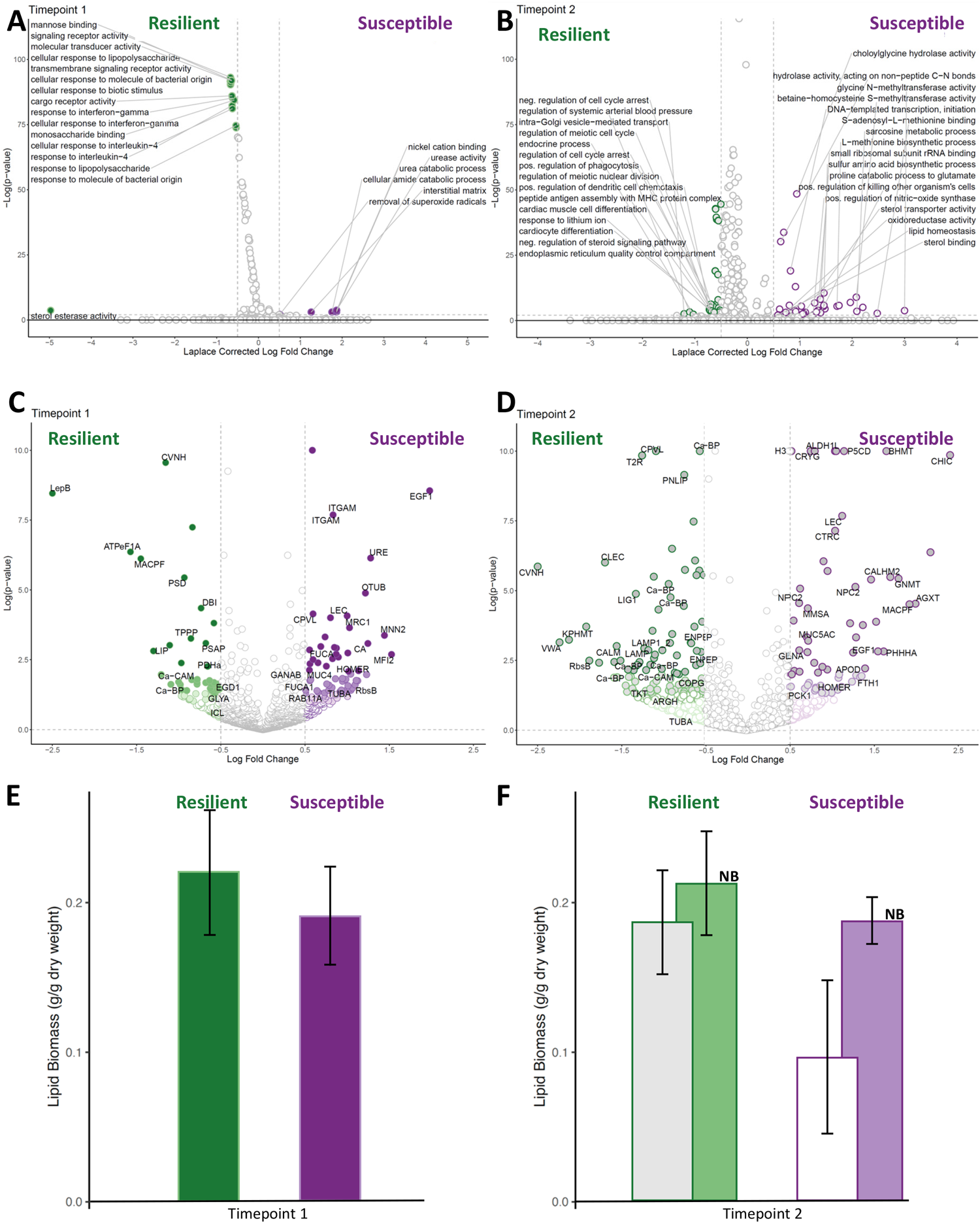
Quantitative molecular data on proteins A-D and lipids E-F, completed on the same 12 colonies used throughout the study (*n=6* susceptible*, n=6* resilient). A and B) Volcano plots depicting -Log (p-value) vs the Laplace corrected Log Fold Change (LFC) for protein-associated Gene Ontology terms. Colored dots signify GO terms that were statistically significantly different between resilient and susceptible corals (Laplace corrected Log fold ≤-0.5 or ≥0.5 and *p-*value ≤0.01). A) T_1_ with negative LFC indicating proteins more abundant in resilient corals (T_1_R) while positive values correspond to proteins that were at higher abundance in susceptible corals (T_1_S). B) T_2_ where negative LFC indicates greater abundance in resilient (T_2_R) *M. capitata* while GO terms with positive values are higher in susceptible corals (T_2_S). C and D). Volcano Plots of individual protein abundances. LFC ≤ −0.5 are proteins that were detected in significantly higher abundance in the resilient coral cohorts (green) at C) T_1_ and D) T_2_. LFC ≥ 0.5 are proteins that were detected in higher abundance in the susceptible coral cohorts (purple) before thermal-stress-induced bleaching (C: T_1_) and after (D: T_2_). E and F) Average lipid biomass (g/g dry weight) measurements on all resistant (green) and susceptible (purple) samples from the different cohorts at E) T_1_ before bleaching and F) T_2_ after bleaching (grey boxes) and non-bleached controls maintained at 25°C (light green and purple “NB” boxes).

### Immune system responses

To elucidate complete metabolic pathways being preferentially utilized by either the resilient or susceptible colonies at the two timepoints, significant differential abundances of proteins were calculated (Fig. 2C, D; significance only reported when *p*≤0.05; Dataset S2G-H). Comparisons of resilient and susceptible colony proteomes before the simulated thermal bleaching event (T_1_), revealed that resilient colonies had 39 proteins at significantly higher abundances than the susceptible cohort, whereas susceptible colonies possessed 56 proteins at significantly higher abundances (Fig. 2C). In T_1_R colonies, signal peptidase (LepB) yielded the highest differential abundance, followed by F-type H+-transporting ATPase subunit alpha (ATPeF1A), and a membrane attack complex component/perforin (MACPF) domain-containing protein known to lyse virus-infected and pathogenic bacterial cells. Susceptible corals before bleaching (T_1_) significantly increased the abundance of Fibropellin-1 (EGF1), a component of the apical lamina, the surface glycoprotein melanoma-associated antigen p97 (MFI2), an enzyme involved in glycosylating proteins, alpha 1,2-mannosyltransferase (KTR1_3), and the urea degrading enzyme urease (URE1).

After the colonies were bleached (T_2_), the resilient cohort was characterized by 108 proteins that significantly increased in abundance while the susceptible cohort significantly increased the abundance of 63 proteins (Fig. 2D). A CyanoVirin-N domain-containing protein (CVNH) was identified to have the most consistent expression across the 12 colonies tested (i.e., lowest *p*-value), a high LFC in the T_1_R proteome, and the highest LFC in T_2_R (Fig. 2D). Further analysis of this protein sequence against the conserved domain database revealed that it contains four CyanoVirin-N conserved domains with viricidal activity that interact with the glycoproteins on the viral envelope (49). Conversely, chitinase (CHIC), an enzyme capable of degrading chitin, exhibited the highest LFC in the susceptible corals post-bleaching (Fig. 2D), another possible indication of symbiont degradation.

Cluster analysis of proteins that were identified across all experiments to be significantly abundant in at least one treatment (LFC ≥|1|, *p*-value<0.01) are represented in a heatmap that spans fifteen metabolic pathways (Fig. 3). Resilient and susceptible coral proteomes are most similar at T_1_ (clusters 1-3, 9-12) compared to the proteomes at T_2_ (Fig. 3). Furthermore, resilient corals prior to bleaching exhibited higher abundances of several proteins involved in anti-viral activity, immune response, and symbiosome maintenance (clusters 4-7) than susceptible corals. After thermal bleaching, the resilient coral colonies maintain a significantly higher abundance of six enzymes, including CNVH (cluster 1), compared to susceptible corals. The T_2_R cohort uniquely increased the abundance of 37 additional proteins (cluster 3) involved in nitrogen metabolism, immune response, endosome/symbiosome activity, and DNA translation, among others. Pre-bleaching susceptible corals (T_1_S), despite exhibiting somewhat similar proteomic trends to T_1_R, revealed one unique cluster (cluster 8) of 9 proteins that were significantly increased in abundance. These proteins play a role in structures and functions such as the extracellular matrix, and immune response, or are associated with lysosome/phagosome activity. Post-bleaching susceptible corals (T_2_S) had the most distinct proteomic response, with depletion of nearly all proteins represented by clusters 1-8 and enrichment in a unique suite of proteins in carbon, nitrogen and lipid metabolism, the biosynthesis of secondary metabolites, the extracellular matrix, and the immune system (clusters 11-12).

**Fig. 3.**
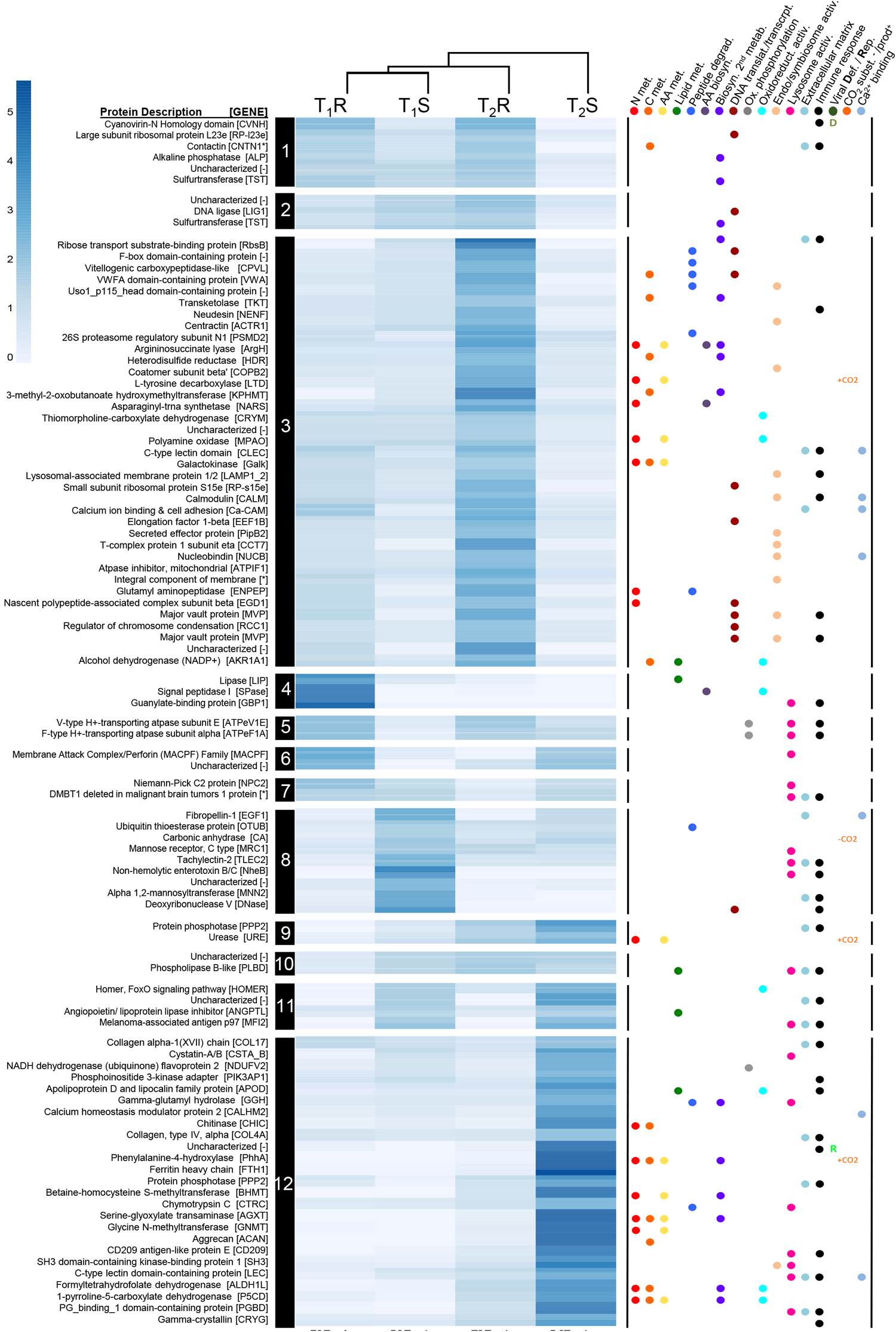
Clustered heatmap of the subset of proteins identified to have a log fold change ≥1 or ≤-1.0 (*p* <0.05) in time point comparisons: T_1_R vs. T_1_S or T_2_R vs. T_2_S. Heatmap shades of blue indicate averaged NSAF values for bioreplicates per condition, normalized by the row mean. Rows are clustered using a correlation algorithm and a dendrogram was set to cut 12 distinct clusters (indicated by #s 1-12 black). Right panel dot-matrix indicates metabolic categories identified through KEGG, UniProt and GO (D) viral defense or (R) reproduction, CO_2_ or as a substrate (CO ^-^) or product (CO ^+^), or Ca-binding domain (i.e., Ca-binding).

### Resilient corals retain lipids through the thermal bleaching event

Previous investigations on recovery from thermal bleaching events revealed that *M. capitata,* unlike other corals, has the unique ability to replenish energy reserves within 1-2 months after the bleaching event, making it one of the coral species with the fastest recovery rates (50). Pre-bleaching, resilient corals had significantly greater abundances of enzymes involved in lipid degradation compared to the susceptible cohort (*e.g.,* PSAP and LIP, Fig. 2A). To determine if pre-bleaching lipid biomass (*i.e.,* T_1_) is a significant and predictable metric to identify *M. capitata* colonies that will recover from thermal bleaching events, total lipids were measured. No significant difference in coral lipid content was found before the simulated thermal event (*i.e.,* between T_1_R and T_1_S colonies; Dataset S1D-E). Coral lipid content varied significantly by bleaching status at T_2_ (T_2_B vs. T_2_NB: *p*=0.00079; Fig. 2F) and long-term tolerance to bleaching (T_2_R vs. T_2_S: *p*=0.00095; Fig. 2E). Interaction of the two variables was also significant (*p*=0.044): susceptible corals experienced a decrease in mean lipid content by 44% after exposure to thermal stress (T_1_S and T_2_S), while resilient colonies decreased by only 16% (T_1_R and T_2_R).

### Resilient corals have more diverse bacterial communities

Alpha diversity of the microbiome based on the 16S rRNA V4 variable region was quantified using Faith’s phylogenetic diversity (Faith’s PD), a measure of microbiome richness that accounts for phylogeny. Faith’s PD significantly differed among groups defined by each combination of timepoint, bleaching resilience, and bleaching status (Kruskal Wallis *p*=0.0045). Prior to bleaching (T_1_), resilient *M. capitata* had higher alpha diversity compared to susceptible corals (Kruskal Wallis *p*=0.016; Fig. 4A). However, this difference could be attributable to multiple comparisons (false discovery rate (FDR) q=0.061). After bleaching, previously low-diversity susceptible corals exhibited significantly increased alpha diversity (*p*=0.0065, FDR q=0.048), while previously high-diversity resilient corals did not (*p*=0.52, FDR q=0.6). Thus, resilient corals showed smaller microbiome changes during bleaching and over time than susceptible corals, consistent with greater stability in microbiome richness.

**Fig. 4.**
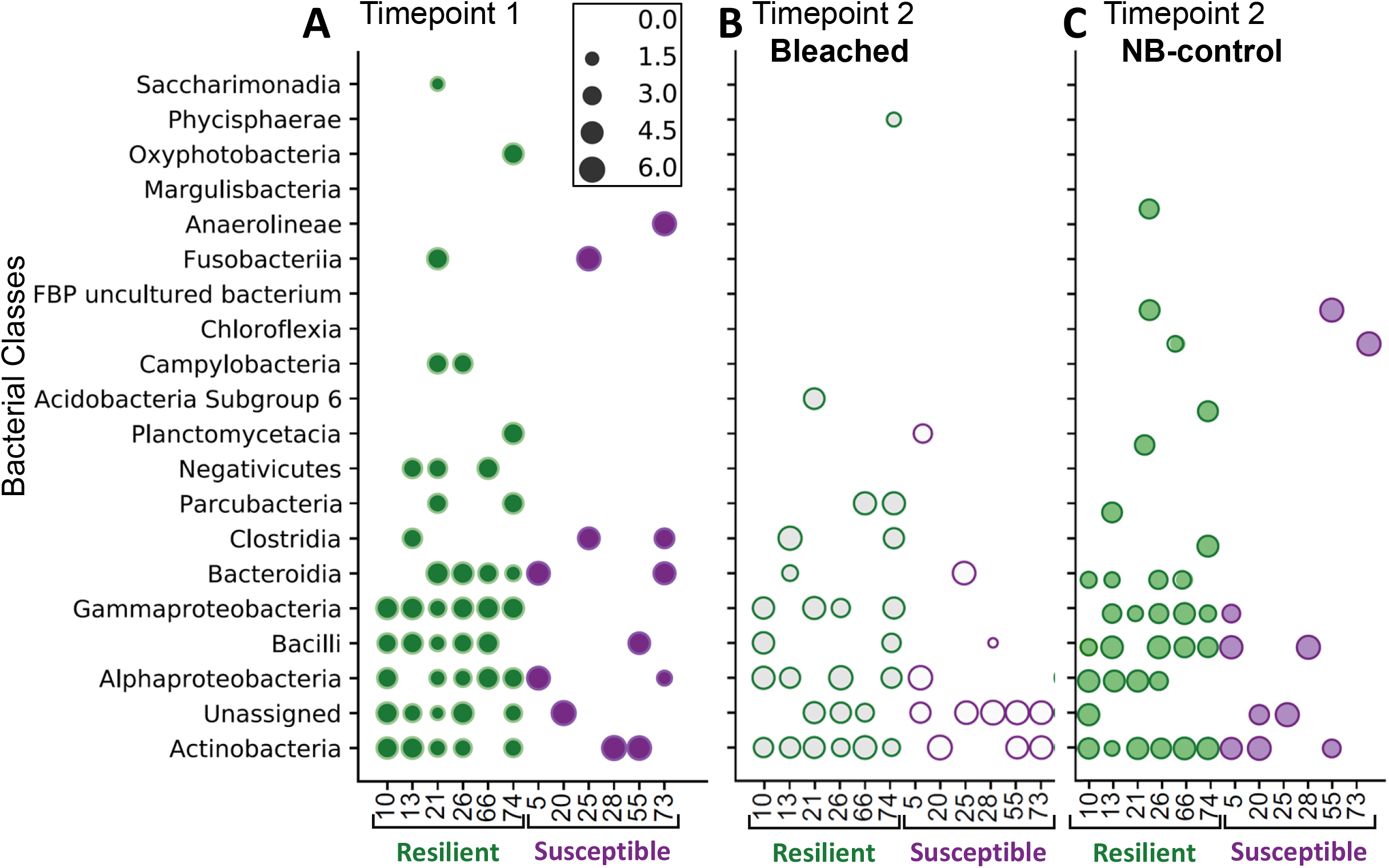
Phylum-level distribution of bacteria identified in resilient and susceptible colonies based on 16S rRNA sequencing data for A) T_1_ (T_1_R: dk green, T_1_S: dk purple), B) T_2_ after thermal stress (T_2_R: grey with green outline, T_1_S: white with dk purple outline), and C) T_2_ control (NB) samples not exposed to thermal stress (NBT_2_R: lt green, NBT_2_S: lt purple). Colony ID numbers are listed on the x-axis. These same 12 colony IDs were used for all analyses presented. Size of the dot represents the log transformation of the phylum-level counts.

Beta diversity was also quantified to determine the similarities of the bacterial communities between cohorts. Across combinations of timepoint, resilience, and bleaching status, microbial community composition differed qualitatively (Unweighted UniFrac PERMANOVA; *p*=0.002) and quantitatively (Weighted UniFrac PERMANOVA; *p*=0.041). These differences were not attributable to differences in microbiome dispersion (Weighted and Unweighted UniFrac *p*>0.05).

The main taxonomic drivers of community differences revealed that Gammaproteobacteria were well represented in the six resilient colonies and nearly absent in susceptible colonies (Fig. 4), consistent with the overall community differences detected in beta-diversity analysis. At the family level, multiple microbial families showed striking differences between the resilient and susceptible cohorts. The clearest of these differences was seen in *Moraxellaceae*, a family in class Gammaproteobacteria that was only found in resilient corals at T_1_. Microbiome Multivariate Association with Linear Models (51), a statistical analysis method that identifies multivariable association between microbial features and metadata (i.e., time, resilience, bleaching), confirmed that *Moraxellaceae* significantly correlated with both time and resilience (MaAsLin2_TIME_ *p*=0.0001; MaAsLin2_RvS_ *p*=0.0007; Table S1). Additionally, the *Caulobacteraceae* microbial family—in the phylum Proteobacteria—was present at elevated abundance in resilient corals, and lower abundance in susceptible ones (irrespective of timepoint or bleaching status (MaAsLin2_RvS_ *p*=0.0006).

## Discussion

Although multiple Cnidarian species have had their proteomes analyzed (*e.g., Eunicea calyculata* (52); *Amphistegina gibbosa* (53); *Acropora microphthalma* (54); *Acropora millepora* (55), *Montipora capitata* (56, 57)), to date, no studies have exhaustively explored pre-bleaching protein-level physiology in combination with multiple other molecular-level factors to determine if there are traits predictive of resilience to bleaching events. Despite previous research showing that symbiont clade D can lead to reduced levels of bleaching in multiple coral species or improve thermal tolerance (29, 36), these coral colonies show no significant differences in clade abundances or distributions across bleaching status, time point collected, or tolerance to bleaching (Dataset S1A, B; Fig S1B). In general, more proteins were detected in all corals at T_2_ post-bleaching relative to T_1_ pre-bleaching, a common response to exogenous stressors (e.g., 58-60). Simultaneous activation of multiple metabolic pathways provides bleached *M. capitata* with new carbon and nitrogen acquisition strategies as symbiont-delivered photosynthate is diminished or absent. Our primary hypothesis is that molecular phenotypic differences - resulting from genetics or epigenetics - will enhance the ability of some individual corals to mitigate the effects of bleaching. Before thermal stress occurred (T_1_), the only significant differences were identified in protein abundances and in the microbiome diversity between susceptible and resilient colonies. Examination of the significantly changing metabolic pathways in the resilient and susceptible cohorts reveals for the first time how nutritional strategies, antiviral mechanisms, and microbiome diversity pre-and post-bleaching dictate survival.

### Phenotypic advantages in resilient corals prior to bleaching

Several proteins in metabolic pathways associated with maintenance of a functional symbiont-host relationship were present at increased abundance in the resilient coral proteome prior to the bleaching event. The primary active pathways that were enhanced in resilient corals pre-bleaching include sterol and lipid degradation, cellular respiration, oxidative phosphorylation, and carbon metabolism (Fig. 2A). Total lipid biomass in the resilient corals had a broader range of values and a higher average than the susceptible cohort (Fig. 2E).

Prior to bleaching, resilient corals utilize heterotrophic feeding and symbiont photosynthate. The GO term sterol esterase was the most significantly increased term in resilient *M. capitata* colonies. Analysis of proteins contributing to this GO term revealed the dominating contributors to the enrichment analyses included a gastric triacylglycerol lipase-like protein (LIP) (Fig. 3; cluster 4) and the lipid-specific degradation enzyme saposin (PSAP; Fig. 2C). These enzymes are typically involved in digestion and likely reside in the coral gastric cavity, suggesting that at T_1_, pre-bleaching, the resilient corals are using a heterotrophic feeding strategy in addition to photosynthate from *Symbiodiniaceae*. This was not unexpected as *M. capitata* has been previously observed to utilize heterotrophy when not bleached (61). T_1_ resilient corals also possessed higher abundance of early endosome antigen 1 (EEA1), an essential protein in symbiosis establishment (i.e., 62-64), providing resilient corals with an advantage for maintaining symbiont relationships compared to the susceptible corals. Utilization of diverse feeding strategies would provide a distinct advantage to the resilient corals as the excess carbon can be shuttled into lipid storage vesicles (61, 65).

Resilient corals present evidence of more active cellular respiration pathways at T_1_ prior to the bleaching event. Although many enzymes in the carbon metabolism pathway are constitutively expressed in resilient and susceptible corals, the increased abundance of isocitrate lyase (ICL; Fig. 2C) provides resilient corals utilization of the glyoxylate shunt, a TCA-cycle bypass that allows cells to complete anabolic reactions with 2-carbon units without losing carbon as CO_2_ (the opposite of what is observed in susceptible corals at T_1_; Fig. S3A). Further, increased abundances of glycine hydroxymethyltransferase (GLYA; Fig. 2C) provides single carbon (1C) units to the cell, fueling the glyoxylate shunt for the generation of larger carbon-storage molecules (*i.e.,* lipids) from the excess 1-2 carbon unit small molecules, again bypassing the generation/loss of CO_2_ (66, 67). Additional evidence, such as increased abundance of pyruvate dehydrogenase (PDHa; Fig. 2C) and multiple acetyl-CoA transferases support increased cellular respiration and the potential to store excess energy in resilient corals.

Prior to bleaching, the resilient corals appear to prime anti-viral activity and exhibited a significantly more diverse microbiome, which likely supported their immune response. Previous demonstrations of “frontloading” immune response or pathogen-fighting enzymes have been shown to increase survival in corals (68). One of the most differentially abundant proteins that was elevated in resistant colonies at T_1_ was a cyanovirin-like protein (CVNH; Figs. 2C), an evolutionarily conserved protein that binds to viruses and blocks entry into the cell (69). CVNH was also detected in higher abundances in the resilient cohort through T_2_, post-bleaching (Figs. 2D and 3, cluster 1). To determine if there were identifiable viral proteins in the whole-coral protein extractions and mass spectrometry analyses conducted, the data was analyzed using a larger database that included 5 coral-associated viral proteomes (Table S2). The number of confident peptides associated with the viral proteins detected were not statistically different between resilient and susceptible colonies at T_1_, yet their presence in whole-holobiont protein extract does corroborate the need for corals to produce antiviral proteins. Additionally, the T_1_R colonies hosted a significantly more diverse microbiome (Fig. 4A), which has a positive effect on host health (70). The resilient microbiome included the *Moraxellaceae* bacterial family, which was only found in resilient coral colonies. *Moraxellaceae* are associated with local wastewater and they are known to have high numbers of antibiotic resistance genes (ARGs) (71, 72). As the coral host’s immune system is activated against pathogenic bacteria and releases antimicrobial defenses, the *Moraxellaceae* bacterial family’s high number of ARGs may provide them with an advantage for long-term residence on the host. As *Moraxellaceae* has been found to be a common component of many shallow water coral microbiomes, these bacteria may be important in shaping a healthy coral holobiont (73). The functional role of this bacterial family’s unique genome in conferring resilience against bleaching to coral colonies is unknown, but the close association of *Moraxellaceae* on resilient *M. capitata* colonies merits further research.

### Molecular signs of stress before thermal induced bleaching in susceptible corals

A detailed proteomics analysis revealed the metabolic processes identified in susceptible corals prior to bleaching, including urea and amide catabolism, nickel binding, and urease activity (Fig. 2A). Redirected nitrogen and carbon uptake pathways in susceptible corals suggest a decrease in symbiont-derived photosynthate at T_1_. Previous molecular-level investigations of symbiont-host relationships have demonstrated that the majority of nitrogen assimilation occurs via the symbiont-directed GS/GOGAT (glutamine synthase/glutamine oxoglutarate aminotransferase) activity or the coral host-directed glutamine synthetase or glutamate dehydrogenase activity (e.g., 74). Prior literature suggests that the majority of nitrogen uptake is from symbiont-transferred metabolites resulting from their utilization of free ammonia in the water column (e.g., 75), although it has been suggested that the assimilation of nitrogen by the host itself is underestimated (76). Here, there is evidence that the susceptible coral colonies utilized urea as their primary nitrogen source at T_1_ (Fig. S3).

We propose that high levels of the urease enzyme may be a biomarker of a dysfunctional metabolic relationship between the coral host and its algal symbiont. Urea, a soluble nitrogen-rich molecule, is degraded intracellularly to yield ammonia and carbon dioxide (CO_2_) via urease enzyme (URE1, Fig. 2A,C and S3). At T_1_ in susceptible corals, GO terms for nickel-binding activity and carbonic anhydrase activity are enriched (Fig. 2A). Proteins associated with these functions provide the required nickel cofactor for urease (77) and increased abundances of carbonic anhydrase enzyme (CA) rapidly converted CO_2_, byproducts of the reaction, into carbonic acid (or bicarbonate). It has been suggested that coral cells rely on this pathway to acquire additional nitrogen when under stress (78, 79). For example, in corals lacking Symbiodiniaceae, urease activity increased to compensate for the lack of Symbiodiniaceae-provided nitrogen (79). Previous experiments on corals revealed that urea- and nickel-enrichments increased photosynthesis and calcification rates, suggesting that these molecules support coral growth in adverse environmental conditions (77). Isocitrate dehydrogenase (ICDH1; FC: 0.57)), an enzyme that is increased under nitrogen starvation, is slightly increased in T_1_ susceptible (compared to T_1_R) corals. ICDH1 links the carbon metabolism (TCA cycle) and nitrogen cycle together to generate glutamate (via GS-GOGAT; Fig. S3A). The increased presence of enzymes involved in these alternate routes of nitrogen and carbon acquisition provide molecular evidence for their use as potential biomarkers of environmental stress and/or the beginnings of dysfunctional symbiosis.

After thermal bleaching (T_2_), the abundance of urease (URE1) continues to increase and is consistently more abundant in susceptible corals at both timepoints (Fig. 3, cluster 9). Therefore, urea-dominated nitrogen acquisition strategy in the host increases as the host-symbiont relationship becomes compromised in susceptible coral colonies responding to thermal stress. Although it has been proposed that the prokaryotic host-associated microbiome could provide bioavailable nitrogen via nitrogen fixation when the host is stressed, the 16S rRNA does not provide species-level resolution that would definitively reveal if any of the noted bacterial families in T_2_S microbiome were nitrogen-fixers.

Importantly, the susceptible corals displayed early evidence for the rejection and degradation of symbiosomes in before the thermal stress starts. The coral’s symbionts reside in specific phagosomes called symbiosomes; corals therefore have specific enzymatic and signaling pathways to disrupt the standard phagosome recycling mechanisms, ensuring the symbiont’s residence. Typical phagosome recycling via hydrolytic enzymes is directed by Rab11 expression in coral hosts (80). Established, healthy symbiotic relationships therefore inhibit the Rab11 pathway, resulting in a decrease in Rab11 abundance (e.g., 80). Susceptible corals at T_1_ displayed significantly increased abundance of Rab11 (Fig. 2C), an early indication of a dysfunctional symbiotic relationship and potential host-directed degradation of the symbiosome, or symbiophagy (81). This host-symbiont disequilibrium hypothesis in T_1_S corals is further supported by increased abundance of Tubulin alpha (TUBA), a phagocytosis protein recognized to be active in symbiont degradation (82), among other functions. Three enzymes detected at significantly increased abundances were involved in glycan degradation, in particular the mechanism involved in cleaving mannose-based oligosaccharides: alpha-L-fucosidase (FUCA1), mannose-receptor (MRC1), and mannosyl-oligosaccharide alpha-1,3-glucosidase (GANAB) (Fig. 3, cluster 8). As mannose is recognized by lectin proteins in corals to identify pathogens and symbionts, the degradation of these mannose-based oligosaccharides would weaken physical associations of the host with symbionts and its microbiome (83, 84). Further, the degradation of these oligosaccharides would specifically provide easily accessed glucose monomers for supporting the energy-demands of the susceptible coral colonies at T_1_. Increased presence of mucin proteins (*i.e.,* MUC4, Fig. 2C), a noted deterrent to pathogen colonization (85), and lectins, pathogen recognition proteins (e.g., TLEC2; Fig. 3, cluster 8) suggest that T_1_ susceptible corals are being challenged by pathogens, further weakening their immune system prior to the bleaching event. The T_1_S microbiome was less diverse than the T_1_R colonies’ microbiomes and more variable across the entire susceptible cohort (*i.e.,* each T_1_S colony had a different taxonomic composition). Quantification of the panel of proteins listed here linked to symbiont rejection could provide coral managers with a rapid biomarker test for identifying which corals are stressed and may not be suitable for propagation, even under optimal environmental conditions.

### Divergent metabolic strategies in resistant and susceptible corals post-bleaching

Constitutive post-bleaching (T_2_) response across all *M. capitata* included many components of the phagocytic and endocytic pathways, indicating that active symbiotic expulsion (86) during thermal stress-induced bleaching was occurring regardless of whether the corals were resilient or susceptible. In particular, Rab11, the inhibitor of symbiosome degradation observed in T_1_S, was detected at increased abundances in both coral groups at T_2_ relative to the T_1_ samples. Other constitutively expressed immune response proteins detected at higher abundances at T_2_ included NOD, MAPK, WNT, and TOLL-like receptors. All of these signaling pathways have been previously observed in corals (87) and their detection suggests that during bleaching the innate immune system is activated. Here, we detected increased abundances of the protein responsible for the irreversible step in gluconeogenesis in susceptible corals, and increased abundances of two irreversible steps in glycolysis in the resilient corals (Fig. S3B). This proteomic evidence suggests that after bleaching, the resilient corals have a more accessible glucose source, whereas the susceptible corals are catabolizing non-carbohydrate sources, such as lipids and proteins. These enzymatic pathway analyses also provide a molecular foundation for the observed 49% decrease in lipid biomass in the susceptible colonies and the insignificant change in lipids in the resilient corals between pre- and post-bleaching (Fig. 2F).

### Resilient corals diversify metabolic pathway utilization to recover from thermal bleaching

After bleaching, several GO terms were enriched in resilient *M. capitata*: amino acid synthesis (methionine, and proline), sulfur amino acid metabolic process, immune response, cell signaling/oxidoreductase, endoplasmic reticulum (ER) organization, oxoacid metabolism, and ribosome assembly (Fig. 2B). Sulfur amino acids, such as methionine, are antioxidants and therefore capable of providing oxidative protection to cells (88). Increased levels of these amino acids in resilient corals could be indicative of increased need for protection against oxidative stress resulting from the heat. Sulfurtransferase enzymes are present in both susceptible and resilient corals, however they are significantly increased in resilient corals (Fig. 3, clusters 1 and 2). Heat stress has been found to also induce an increase in endoplasmic reticulum transcripts in *Acropora hyacinthus* (89), mirroring our findings of increased abundance of ER proteins. Increases in cell signaling and ribosome assembly proteins are likely indicative of more normal cellular trafficking in healthy host tissue, enabling recovery from thermal bleaching. Several metabolic pathways are discussed below that support the resilient metabolism through thermal stress compared to the susceptible coral cohort.

Resilient corals activate endocytic uptake pathways and heterotrophic feeding after bleaching event to aid in nutritional recovery. Multiple enzymes involved in endocytosis are increased in the T_2_R cohort providing a heterotrophic avenue for carbon and nitrogen acquisition (Fig. 3, cluster 3). Increased abundance of 2 tubulin alpha proteins (TUBA), vacuolar sorting endocytic protein (VPS4), dynamines and coatamers (COPG, COPB2) imply increased endocytic activity of particles, such as pathogens or food (82, 90). Significantly increased abundance of lysosome associated membrane protein (LAMP) may indicate that symbiont engulfment and degradation is an additional potential source of nutrition for resilient corals post bleaching (81). Significantly increased peptide degradation enzymes in the T_2_R cohort included glutamyl amino peptidase (ENPEP, (Fig. 3, cluster 3), which cleaves acidic amino acids from the N-terminus of peptides for subsequent degradation to enhance cellular growth. T_2_R also significantly increased a vitellogenic carboxypeptidase (CPVL), a protein involved in the degradation of yolk proteins. This may be a sign of a physiological switch to sacrifice reproductive potential to increase the chances of bleaching recovery and short-term survival. Abundant protease/peptidases (Fig. 3, cluster 3) and lipases (*e.g*., triacylglycerol lipase PNLIP; Fig. 2D) in resilient colonies can break the bonds of macromolecular complexes to generate mobile small molecules that can be recycled or further degraded for energy. In support of these findings suggesting adequate nutritional resources were available in the resilient cohort post-bleaching, the increased abundance of transketolase (TKT) in T_2_R may indicate a higher abundance of thiamine (vitamin B1) compared to T_2_S (91, 92).

To further aid in recovery, resistant corals post bleaching appear to utilize several new pathways to aid in cellular nitrogen and carbon demands. Although the urea degrading enzyme URE1 is detected in both T_2_S and T_2_R corals, T_2_R displayed increased abundance of polyamine oxidase (MPAO; Fig. 3 cluster 3) which may allow resilient colonies to access polyamine-nitrogen as needed and produce beta-alanine. Beta-alanine (aminoproponoic acid) is a degradation product of the nucleotide uracil and is a precursor to acetyl-CoA. Notably, it has been found to increase cellular oxygen consumption and respiration rates (93). Detecting multiple enzymes involved in these pathways to be at significantly higher abundance in the T_2_R colonies provides a molecular explanation for hypothesized improved energy production compared to the T_2_S colonies when symbiont derived photosynthate is diminished. Further, this energy may have provided resilient colonies the ability to significantly increase multiple enzymes responsible for DNA transcription and translational processes (Fig. 3, cluster 3).

T_2_R also launched an antiviral campaign during thermal stress to assist the immune system. Cyanovirin protein (CVNH) was detected in resilient corals at T_1_ and T_2_ at three-fold higher abundance, compared to susceptible colonies (Fig. 3, cluster 1). Although we do not believe this to be the only mode of protection for the resilient cohort, high production of this protein could increase the resilience of these corals after bleaching events when they are simultaneoulsy coping with multiple stresses.

### Catastrophic metabolic choices in T_2_S corals

GO term enrichment analysis revealed a greater abundance of proteins participating in peptide degradation and protein transport in susceptible *M. capitata* post-bleaching (Fig. 2B,D; Fig. 3 cluster 12). Since Symbiont-derived photosynthate nutrition is absent in the bleached corals, susceptible colonies may have increased mobilization and degradation of proteins and peptides to provide the needed energy for cellular maintenance. Decreases in the free amino acids pool resulting from protein degradation in thermally bleached *Acropora aspera* suggests that these amino acids are being metabolically leveraged to provide energy during low photosynthate yield (94).

Specifically, susceptible hosts post-bleaching express an abundance of enzymes that suggest host-directed catabolism of remaining symbionts. Several lysosomal-targeted peptidases/degradation enzymes were significantly increased in the susceptible corals in response to bleaching (e.g., cathepsin CATL, galactosidase GLB, and Niemann Pick C2 protein NPC2, Fig. 3, clusters 7, 12). The early signs of a weakened host-symbiont relationship for T_1_S corals (discussed earlier) appears to have progressed further in the susceptible cohort by T_2_. At T_2,_ health-compromised/dead symbionts may be leaking organic substrates that are degraded by lysosomal and intracellular peptidases and hydrolase enzymes (Fig. 3, cluster 12). NPC2 enzymes are concentrated in the symbiosome and participate in the direct sterol transfer from symbionts (95). Evidence of increased host-directed transfer of sterols from the symbiont combined with increased lysosomal-catabolic processes indicate that after bleaching, the symbiosome and its contents are targeted for rapid degradation in susceptible colonies (81).

Further, the decrease in symbiont-derived photosynthate in susceptible corals leads to activated gluconeogenesis and the degradation of glycine betaine. Glucose is one of the primary carbon molecules transferred to holobionts in cnidarian dinoflagellate symbiosis (96, 97). The increased abundance of the irreversible enzymes pyruvate carboxylase (PC) and phosphoenolpyruvate carboxykinase (PCK1) in susceptible corals reveals that gluconeogenesis (*i.e*., the generation of glucose from pyruvate) is more active than glycolysis (*i.e*., the degradation of glucose; Fig. S3B). Increased abundance of these enzymes suggests that susceptible corals are more glucose-limited compared to the resilient corals after bleaching events. Gluconeogenesis depends on the catabolism of amino acids, glycine betaine, and lipids (Fig. 3, cluster 12). Lipid degradation is evidenced by the significant decrease in total lipid biomass between T_1_S and T_2_S colonies (Fig. 2E-F). Only recently were glycine betaines recognized to be a major reservoir of nitrogen for corals and the near-complete glycine betaine catabolic and biosynthesis pathways have been uncovered in several genomes of cnidarians (refs within:, 98). T_2_S corals increased abundance of the betaine-degrading enzymes betaine-homocysteine S-methyltransferase (BHMT), glycine N-methyltransferase (GNMT), and sarcosine dehydrogenase (SARDH). Ngugi et al., (98) suggest that glycine betaines are abundant nitrogen reservoirs that are easily degraded into other nitrogen compounds such as amino acids.

Susceptible coral proteomes post bleaching also revealed a trend of potential decreased immune function at T_2_. Immune pathways results indicate that the NOD, MAPK, and TOLL-like signaling pathways are suppressed in T_2_S corals (*p*<0.10; Dataset S2H). The suppression or inactivation of these important immune pathways may make the corals vulnerable to disease and colony mortality. The suppressed beta diversity in T_2_S corals reveals a shift to a less diverse symbiotic bacterial community, which may be an indicator of the onset of infection (refs within:, 99). Susceptible *M. capitata* colonies also increased tyrosinase (TYR), an indicator of immune response to an infection (100) or neutralization of reactive oxygen species (101) demonstrating that T_2_S corals are being challenged.

### Coral management and restoration applications

As the ultimate goal for coral management is to be able to predict resilient coral phenotypes before investing time and money in restoration, a rapid assay to determine health status is needed. Here we presented three significant differences in the resilient and susceptible coral colonies before the thermal bleaching to forecast long-term health through thermal events: proteins (Fig 2A,C), lipids (Fig. 2E), and microbiome diversity (Fig. 4A). Previous work on corals has revealed that decreases in lipid content and in microbiome diversity can be associated with a range of environmental responses and are not exclusively associated with susceptibility to bleaching stress. Here we propose a protein-based assay to predict resilience and capture more informative results on the molecular-level health of *M. capitata.* We have identified seven proteins that could be quantified in corals before bleaching events as a resilience-based assay to select colonies for propagation or other management strategies (Fig. 5). If using mass spectrometry, quantifying five peptides through the detection of ≥5 diagnostic fragment ions from each of these proteins would provide the user with high confidence on both positive and negative signals of pre-bleaching resilience. This short list could also be expanded to generate a 60 minute assay with up to 250 peptides that are simultaneously monitored, providing further information on heterotrophic feeding/lipid degradation (*i.e*., LIP, PSAP), antiviral activity (*i.e.,* CNVH), symbiophagy (*i.e.,* RAB11, TUBA), pathogen recognition (*i.e*., TLEC), mucin proteins (*i.e.,*MUC4), urea degradation (*i.e.,*URE1), and mannose degradation (*i.e.,* FUCA1, MRC1, GANAB). Alternatively, selected proteins identified here as resilience biomarkers could be developed into a hand-held rapid antigen test with multiple test and control lines that could be assessed in the field on rice-grained size coral tissue samples.

**Fig. 5.**
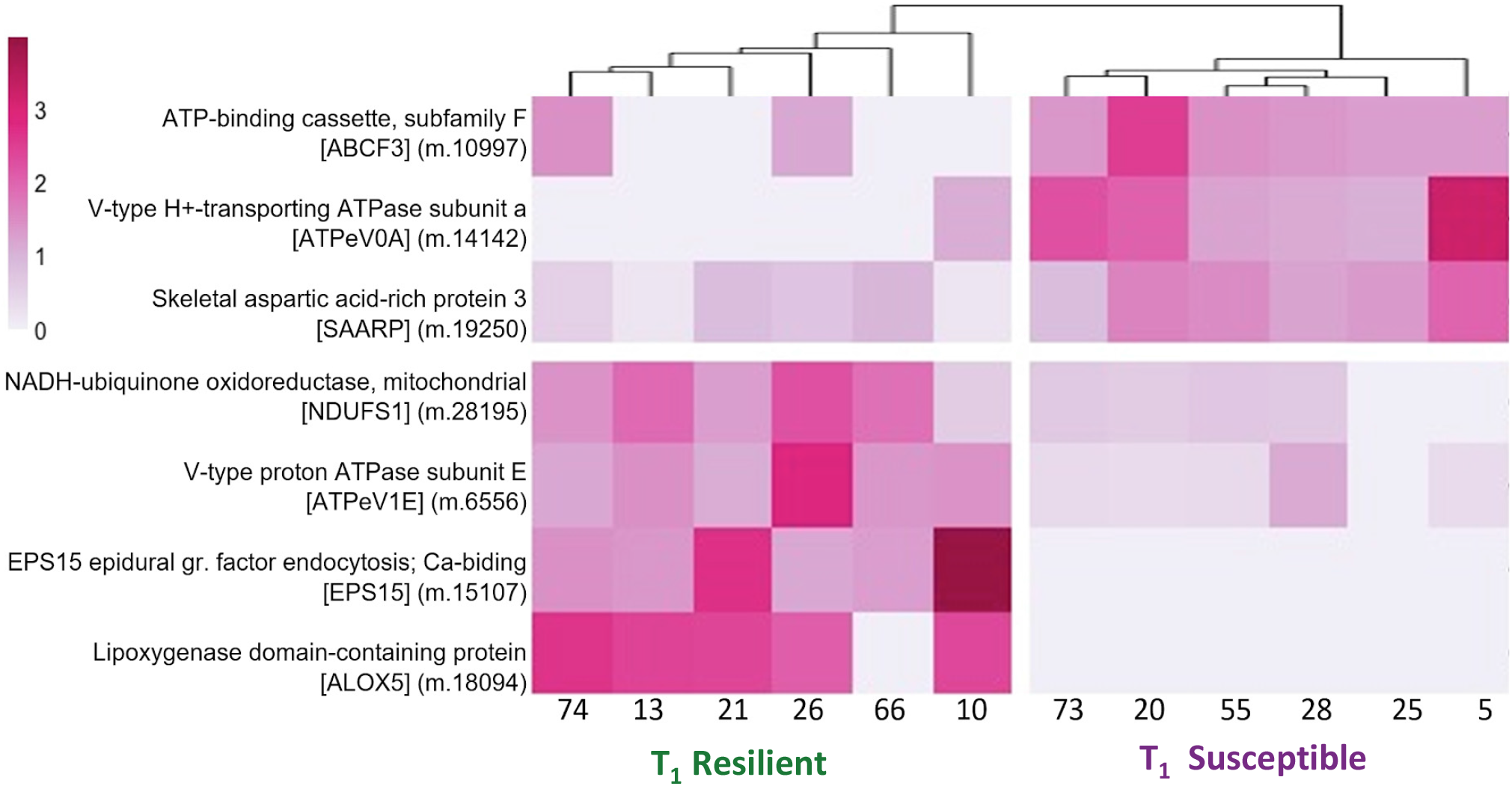
Heatmap of significantly different proteins (p<.01) identified between resilient and susceptible corals at T_1_, pre-bleaching event. Colors depict NSAF values for each of the twelve coral bioreplicates, normalized by row mean. Clustered dendrograms were completed with the correlation algorithm on the x and y axis to generate groups for significantly increased or decreased abundances in the resilient vs. susceptible cohorts.

### Concluding Remarks

This is the first study to use an analytical toolbox that included mass spectrometry-based proteomics, 16S rRNA analyses of the microbiome, total lipids, and symbiont density and diversity to identify intrinsic differences that confer recovery and survival in corals before thermal bleaching events. This study is also unique in that the *Montipora capitata* coral colonies had unexpected, yet vastly different, outcomes from the thermal bleaching event despite identical environmental histories and, to the best of our knowledge, coral genotypes. Our intent was to reveal a multi-factor molecular-level approach for confidently identifying resilient and susceptible coral colonies so that environmental managers could rapidly select quality candidates for propagation while in the field. Despite monitoring a range of physiological and molecular metrics, very few significant differences were discovered in the study that could positively identify resilient colonies before a thermal bleaching event. Despite lipids being a tractable metric in the field, the differences in the resilient vs. susceptible coral lipids were only present *after* the thermal bleaching event, making lipids ineligible as a biomarker for long term survival *prior* to thermal stress. Additionally, lipids are non-specific biomarkers since their levels are often influenced by a variety of physiological factors (*i.e.,* infection, reproduction, tissue thickness, etc.). Promising and distinct differences observed in the 16S rRNA analysis of bacterial diversity revealed that the resilient cohort hosted a significantly more diverse microbiome before the thermal event. Although microbiome diversity could aid as a metric for selecting healthy, robust coral colonies, microbiome stability and diversity can be linked to water chemistry (102), temperature (14), and short or long term diseases (103, 104). Additional research needs to be conducted to determine if specific microbial clades are significantly correlated to coral recovery and resilience through thermal induced bleaching and what functional roles they play. Quantitative proteomics analyses identified proteins that were significantly different in the two cohorts before the bleaching event, that could 1) allow confident predictions in selection of resilient over susceptible colonies and 2) reveal specific molecular advantages in the form of active pathways and primed immune responses that allow resilient *M. capitata* corals to survive the thermal stress despite the expulsion of *Symbiodiniaceae.* Resilient corals have a significantly higher abundance of antiviral proteins and express multiple enzymes involved in a diverse range of carbon and nitrogen acquisition such as lipid degradation, heterotrophic feeding, and respiration. Conversely, colonies that did not survive thermal bleaching had pre-bleaching molecular markers at elevated abundances that play an active role in symbiont rejection, pathogen recognition, and mannose and urea degradation. The proteins represented in each of these pathways and cellular mechanisms can be fully developed into rapid molecular assays to help assess corals and guide mitigation strategies deployed by reef management.

## METHODS

### Coral Colony Collection and Experiment

Seventy-four coral colonies of *Montipora capitata* (approximately 30 cm in diameter) were collected from Moku O Lo’e island (patch reef) located in Kāne’ohe Bay, O’ahu, Hawai’i (21.428°N, 157.792°W) in August 2017. Colonies were brought to shore and acclimated in flow-through outdoor tanks at the Hawaii Institute of Marine Biology (HIMB) for two weeks. At the time of collection, colonies were divided in two pieces to compare physiological performance for the same genotypes with/without exposure to thermal stress (Fig. 1). In September, one half of each coral colony was exposed to warmer water temperatures to simulate a natural thermal bleaching event (56). To reach the 30°C temperature goal for the bleaching treatment, experimental tank temperatures were increased 2°C per day (1°C every 12 hours) for four days. The colonies were rotated once a week between tanks to minimize tank effects. For the bleaching experiment, *M. capitata* colonies were kept at this elevated temperature for three weeks to induce complete coral bleaching in all the colonies. After bleaching occurred, the tank temperature was lowered, following the previously described rate, back to ambient temperature (22°C) and subsamples of coral were taken. The coral halves that were not exposed to thermal stress remained at 25°C and were also rotated within the tank to minimize tank effects. Then, all corals (bleached and not bleached) were placed on racks off HIMB to monitor survival and physiological recovery *in situ* for eight months. Bleaching assessments were conducted on all colonies every week using the Coral Watch Card (The University of Queensland, Australia), along with assessments of mortality (Fig. S1A). Colonies that bleached and recovered were deemed to be part of the resilient cohort while colonies that bleached and died were deemed susceptible to bleaching. All resulting proteomic search results, protein accession numbers and annotation files, lipid data, symbiont density and clade data, chlorophyll data and R code for plot generation and data analysis have been deposited in GitHub (https://github.com/Nunn-Lab/Publication-coral-resilience).

Branches from twelve *M. capitata* colonies were collected at two time points: 1) in September after temperature acclimation in the tanks but before colonies were bleached (T_1_) and 2) in late September, 24 hours after bleached colonies were gradually returned to ambient temperature (T_2_). All coral samples were collected 1 cm from the tip of a branch and snap frozen immediately in liquid nitrogen. Frozen samples were shipped to the University of Washington on dry ice and stored at −80°C. Samples for protein extraction consisted of 2 mm thin cross-sections of the branches, encompassing both tissue and skeletal matrix.

### Symbiont and Chlorophyll analyses

Chlorophyll a concentrations and dinoflagellate symbiont (Symbiodiniaceae) densities from each of the colonies were investigated (Dataset S1A). Briefly, chlorophyll a was extracted with 100% acetone and absorbance was measured with a light spectrophotometer (Dataset S1B). Symbionts were separated from triplicate ground coral tissue by centrifugation and symbiont pellets were homogenized prior to being counted using a hemocytometer. Chlorophyll a and symbiont densities were standardized to grams of ash-free dry weight (gdw) of coral tissue (Dataset S1A,C). In order to assess the ratio of Symbiodiniaceae C and D clades a 4mm piece of frozen *Montipora capitata* was crushed using a frozen mortar and pestle and total DNA was extracted and Quantitative Real Time PCR (qPCR) assay of the symbionts’ actin genes was used to determine the ratio of Symbiodiniaceae C and D clades (Dataset S1B, Fig. S1B). Further details found in SI Methods.

### Lipid Analyses

Total lipids were analyzed on each sample following the methods of Rodrigues and Grottoli (50). Briefly, whole fragments (tissue plus skeleton) were crushed and digested in a 2:1 chloroform:methanol solution, sequentially washed in a 0.88% KCl solution, dried under grade 5.0 N_2_ gas to a constant weight (Dataset S1D-E)

### Proteomics

Six colonies from the resilient cohort and six colonies from the susceptible cohort were randomly selected as bioreplicates to track phenotypic differences in protein abundance through time (*i.e.,* T_1_ and T_2_). Details can be found in SI Methods. Briefly, proteins were extracted from whole coral fragments (4 mm diameter x 1 mm thick, tissue plus skeletal matrix) and resulting protein concetrations were determined with bicinchoninic acid (BCA) Protein microplate assay. Protein lysates (50µg per coral sample) were reduced, alkylated, and digested Trypsin (modified porcine sequencing grade trypsin; Promega; 1:20 enzyme:coral protein). Each digested peptide sample was amended with Peptide Retention Time Calibration Mixture (PRTC; Pierce) such that 50 fmol of PRTC was analyzed with 1 µg of coral peptides for each mass spectrometry experiment.

*M. capitata* samples were analyzed using liquid chromatography coupled to tandem mass spectrometry (LC‒MS/MS) on a Q‒Exactive‒HF (Thermo Scientific) in Data Dependent Acquisition (DDA) Top 20 mode. Samples were separated using a heated (50°C) 40 cm long analytical column packed with C18 beads (Dr. Maisch HPLC, Germany, 0.3 µm, 120Å). Peptides were chromatographically separated on a Waters nanoAcquity UPLC using an acidified (0.01% formic acid) acetonitrile:water gradient of 2–45% over 120 minutes. Internal and external standards were monitored to ensure peptide peak area correlation variances were <10% through the duration of the analyses. Data was searched against a translated *M. capitata* transcriptome (105) GSE97888_Montiporacapitata_transcriptome.fasta). Protein identifications from the whole-cell lysates are reported if two or more peptides were identified, at least one terminus was tryptic, and the false discovery rate <0.01) (Dataset S2A-E). Differential relative protein abundances for resilient vs. susceptible corals were determined for each timepoint (T_1_ and T_2_) using the QPROT-QSPEC package (106)(Dataset S2G-H). Differential abundances of proteins are reported with the following *p*-value cutoff rules: 1) *p*<0.10 if several proteins within a pathway, 2) *p*<0.05 if significance of an individual protein, or 3) *p*<0.01 if identifying a potential biomarker.

### MetaGOmics Biological Enrichment Analysis

To determine if categories of proteins were enriched in the resilient vs. susceptible coral cohorts at the two timepoints, a biological enrichment strategy that analyzes Gene Ontology (GO) categorical terms was used to compare sets of detected proteins (48). Top results are reported with a cutoff E-value <1E-10. A fasta file of all *M. capitata* protein sequences confidently identified in these experiments (File S3) was analyzed with MetaGOmics v.0.1.1. Although MetaGOmics was designed to analyze microbiomes, the use of the software was modified to work with a single organism by ignoring the taxonomic enrichment analysis to instead examine functions that are significantly enriched or depleted in pairwise comparisons of coral cohorts. Additional details found in SI Methods.

### Microbiome 16S rRNA Analyses

Total DNA was extracted from the corals selected for this study (n=12) using the Qiagen DNA extraction kit. All 16S rRNA gene amplicon sequence data, processing steps and code for quality control on the microbiome data and analysis are available on GitHub (https://github.com/tanyabrown9/Resilient_vs_Susceptible_Mcapitata). Sequences are deposited in NCBI as bioproject PRJNA933787. Initial sequencing resulted in a collection of 2,472,819 total reads, with an average read depth of 68,689 (± 28,107 SD) sequences per sample (Dataset S4A-B). Amplicon sequence data were processed using the QIIME2 software package (107). Alpha diversity was assessed using the number of unique observed ASVs in rarefied samples by the Simpson’s Evenness and Shannon’s Diversity Indexes. Overall differences in alpha diversity across susceptibility and time points were tested using Kruskal-Wallis tests. Post-hoc comparisons were performed within each group as well as combined comparisons with *p*-values for pairwise tests between treatments adjusted for multiple comparisons using Bonferroni correction. Beta diversity was assessed between samples using Weighted UniFrac distances and Bray-Curtis dissimilarities. The significance of differences in beta-diversity between susceptibility and time was tested using PERMANOVA (108). The top 10 bacterial families in each sample type were selected for taxonomic analysis. Significant differences between bacterial families, susceptibility, and timepoint were carried out using a nested ANOVA. Multivariate Association with Linear Models was performed on the 16S data using the R package MaAsLin2 (51). Additional details can be found in SI Methods.

## Supplemental Information

## Data, Materials, and Software Availability

All raw MS proteomic data and protein FASTA files used for searching can be accessed at PRIDE accession PXD021262 (UsernameXXXXX). All resulting proteomic search results, protein accession numbers and annotation files, lipid data, symbiont density and clade data, chlorophyll data, and R code for plot generation and data analysis have been deposited in GitHub (https://github.com/Nunn-Lab/Publication-coral-resilience). All 16S rRNA gene amplicon sequence data, processing steps, and code for quality control on the microbiome data and analysis are available on GitHub as well (https://github.com/tanyabrown9/Resilient_vs_Susceptible_Mcapitata). All study data are included in the article and/or as SI Datasets. All sequences used in this study are publicly available through NCBI GenBank and ProteomeXchange PRIDE. Accession numbers and annotations are provided as supplementary files (Datasets S1-3).

## Acknowledgments

We would like to thank the Gates Coral Lab for hosting and supporting us during this experiment at the Hawaiʻi Institute of Marine Biology. We thank Brenner Wakayama, Gavin Kreitman, Melissa Jaffe, and Sean Frangos for their help with collecting and culturing the experimental corals. Work was supported in part by the University of Washington’s Proteomics Resource (UWPR95794), by NSF IOS-IEP 1655682 awarded to B.L.N. and J.L.P.G., NSF IOS-IEP 1655888 to L.J.R., NSF GFRP awarded to J.B.A., NIH-F31 awarded to M.M and Alfred P. Sloan Research Fellowship to J.L.P.G. and by C.C.N (private funds to support environmentally relevant research aimed at making a difference) to B.L.N.

