## Supplemental Information for "Resilience in a time of stress: revealing the molecular underpinnings of coral survival following thermal bleaching events"

**This PDF file includes:**

- Supporting text
- Figures S1 to S4
- Tables S1 to S2
- Legends for Datasets S1 to S4
- Legends for Files S1 to S3
- SI References

**Other supporting materials for this manuscript include the following:**

- Datasets S1 to S4
- Files S1 to S3

### Supplemental Methods

**Symbiont and Chlorophyll analyses.** Chlorophyll a concentrations and dinoflagellate symbiont (Symbiodiniaceae) densities from each of the colonies were investigated (Dataset S1A). Briefly, chlorophyll a was extracted with 100% acetone and absorbance was measured with a light spectrophotometer (Dataset S1B). Symbionts were separated from triplicate ground coral tissue by centrifugation and symbiont pellets were homogenized prior to being counted using a hemocytometer. Chlorophyll a and symbiont densities were standardized to grams of ash-free dry weight (gdw) of coral tissue (Dataset S1A,C). In order to assess the ratio of Symbiodiniaceae C and D clades a 4mm piece of frozen *Montipora capitata* was crushed using a frozen mortar and pestle and total DNA was extracted using the Qiagen DNeasy Plant Mini Kit according to manufacturer's protocols. DNA concentrations were assessed using the NanoDrop 2000. A Quantitative Real Time PCR (qPCR) assay of the symbionts' actin genes (Cunning & Baker 2013) was used to determine the ratio of Symbiodiniaceae C and D clades (Dataset S1B, Fig. S1B). M. capitata host DNA was quantified using the PaxC intron (Cunning, Ritson-Williams, and Gates 2016). All samples (both clades and host DNA) were run in triplicate on a BioRad CFX 96 qPCR thermalcycler. Each 23 µl reaction contained 3.68µl SYBER Green master mix (BioRad), 0.2µM each primer and 1.5mM MgCl2. PCR cycles were conducted as follows: 95°C for 3min followed by 40 cycles of 95°C for 20 sec, 50°C for 20 sec and 72°C for 20 sec. This was followed by a denaturization curve to confirm only one product was amplified. A two-way ANOVA was applied to examine the impacts of timepoint, bleaching resilience, and bleaching status on percentage of clade C and D in the host tissue.

**Proteomics.** Six colonies from the resilient cohort (colony numbers R.10, R.13, R.21, R.26, R.66, R.74) and six colonies from the susceptible cohort (colony numbers S.5, S.20, S.25, S.28, S.55, S.73) were randomly selected as bioreplicates to track phenotypic differences in protein abundance through time (*i.e.,* $T_1$ and $T_2$). Proteomic analyses were not completed on $T_2$ control colonies that remained at 25°C unbleached. A total of 24 samples of frozen coral (4 mm diameter x 1 mm thick, tissue plus skeletal matrix) were manually smashed using a mini mortar and pestle and resulting tissue and debris were sonicated (5 x 10 s) in 6 M urea in 50 mM $NH_4HCO_3$. Sample tubes were cooled between each sonication in a bath of ethanol and dry ice (30 s). Protein concentrations were measured using the Pierce BCA Protein Kit microplate assay (Thermo Scientific) following the manufacturer's protocol.

Protein digestions were conducted with 50 µg of protein per coral sample. Protein lysates were solubilized in 100 µl of 6 M urea buffer, reduced with 500 mM TCEP (1 µl), and pH was controlled with 1.5 M Tris pH 8.8 (6.6 µl) and adjusted as needed to pH > 8. Proteins were alkyated with iodoacetamide (IAA; 200 mM, 20 µl) and remaining alkylating agent was quenched with diothiothreitol (DTT; 200 mM, 20 µl). Prior to trypsin digestions, the urea was diluted with 25 mM $NH_4HCO_3$ (800 µl) and HPLC grade methanol (200 µl). Trypsin (modified porcine sequencing grade trypsin; Promega; 1:20 enzyme:coral protein) was added to enzymatically cleave proteins overnight at room temperature. Sample pH was then modified to $\leq$2 with 10% trifluoracetic acid (TFA) and peptides were evaporated to dryness at 4°C in a speedvac. Dried coral peptide samples were reconstituted in 5% acetonitrile and 0.1% trifluoroacetic acid and desalted on C18 macrospin columns (The NestGroup) following the manufacturer's guidelines. Eluted peptides were dried to final volume <10µl on a speedvac and reconstituted in 200 µl 2% acetonitrile and 0.1% formic acid. Peptide Retention Time Calibration Mixture (PRTC; Pierce) was added to each coral peptide sample as an internal standard to ensure consistency of peptide detection. Each sample was amended with PRTC such that 50 fmol of PRTC was analyzed with 1 µg of coral peptides for each mass spectrometry experiment.

*M. capitata* samples were analyzed using liquid chromatography coupled to tandem mass spectrometry (LC–MS/MS) on a Q–Exactive–HF (Thermo Scientific) in Data Dependent Acquisition (DDA) mode. An in–house analytical column (40 cm long) was packed with C18 beads (Dr. Maisch HPLC, Germany, 0.3 µm) and heated (50°C) during analysis to maintain steady backpressure. Peptides were chromatographically separated on a Waters nanoAcquity UPLC using an acidified (0.01% formic acid) acetonitrile:water gradient of 2–45% over 120 minutes. MS1 was collected on 400-1200 m/z with a 70,000 resolution and AGC target of 1e6; MS2 were collected with a loop count of 20 excluding +1 and >+6 MS1 ions using a 10 s dynamic exclusion, 35,000 resolution, and AGC target of 5e4. Sample analyses were randomized, and quality controls were analyzed every 11th injection. Select peptides from QC samples were monitored using Skyline (1) to ensure that peptide peak area correlation variances were <10% through the duration of the analyses. Three bioreplicates of susceptible corals did not yield high quality mass spectrometry results due to extensive intracellular degradation (*i.e.,* cell- induced death) and were not utilized in any of the analyses ($T_2$ samples S.20, S.25, S.28). The raw MS data is at PRIDE accession PXD021262 (Username:; Password: r5h4H1vo)

From each mass spectrometry experiment *M. capitata* peptides were identified and proteins were inferred using a proteome database derived from the *M. capitata* transcriptome (2) GSE97888_Montiporacapitata_transcriptome.fasta). The transcriptome was translated using Transdecoder v 2.0.1 (3). This final *M. capitata* proteome was concatenated with 50 common contaminants (cRAPome) and the internal standard PRTC peptides to generate our search database (File S1). The raw MS data were searched against the protein database using Comet v 2016.01 rev.3 (4)). Comet parameters included concatenated decoy search, 20 ppm peptide mass tolerance, enzyme set at trypsin with two missed cleavages allowed, variable cysteine modification of 57 Da, and fixed methionine modification of 15.999 Da. Concatenated target–decoy databases searches were completed and minimum protein and peptide thresholds were set at $P$> 0.95 on ProteinProphet and PeptideProphet. Protein identifications from the whole-cell lysates were accepted by ProteinProphet if the above-mentioned thresholds were passed, two or more peptides were identified, and at least one terminus was tryptic (false discovery rate <0.01) (5). Resulting data files across all samples were analyzed with Abacus (6) to generate consistent protein inferences across replicates (*i.e.,* joining homologous protein identifications if needed) and calculating normalized spectral abundance factors (Dataset S2I). Proteins were considered high confidence if two unique peptides were identified across all mass spectrometry experiments. For each treatment and time, a subset from that list was generated and included all proteins whose sum of the total spectral counts was >2 for the specific cohort (Dataset S2A-D and E).

Differential relative protein abundances for resilient vs. susceptible corals were determined for each timepoint ($T_1$ and $T_2$) using the QPROT-QSPEC package (7). QPROT-QSPEC package first normalizes each dataset and then calculates the posterior distribution of the log fold change and the associated Z-statistic (6, 7)(Dataset S2G-H). The Z-test is similar to the T-test but accommodates for missingness of data when looking at sample size comparisons to determine significance levels to report or compare. Z-scores were converted to $p$-values using R (pvalue2sided=2*pnorm(-abs(z)) to simplify reporting. Differential abundances of proteins are reported with the following *p*-value cutoff rules: 1) $p<0.10$ if several proteins within a pathway, 2) $p<0.05$ if significance of an individual protein, or 3) $p<0.01$ if identifying a potential biomarker.

Volcano plots were made with resulting QPROT statistics in R ggplot v 3.0.4 (8), and the heatmaps were generated with pheatmap v. 1.0.12 (9). Heatmap (Fig. 3) was generated by averaging NSAF values for each sample suite (*i.e.,* $T_1S$, $T_1R$, $T_2S$, $T_2R$) for a defined set of proteins that meet the established criteria (LFC $\geq|1|$, *p*-value<0.01) while the biomarker heatmap (Fig. 5) was generated from NSAF values for each bioreplicate to demonstrate quantitative consistency. Vertical clusters were completed with the "correlation" algorithm.

All proteins identified were analyzed with BlastKOALA (10) to find associated K numbers from the Kyoto Encyclopedia of Genes and Genomes (KEGG) and a list of metabolic pathways in which the enzymes are putatively involved. The top-scoring K numbers, KEGG descriptions, and gene names are reported. The *M. capitata* proteome was also analyzed with BLASTp against UniProtKB Swiss-Prot non-redundant protein database (11) (downloaded 10.15.2018) using the diamond algorithm (12). Names of proteins in the manuscript text and figures are reported using the following hierarchy: 1) KEGG (gene name, description, metabolic pathways), 2) Uniprot BLAST (gene name, description), and 3) Uniprot BLAST Gene Ontology (GO Biological Process) (Dataset S2F). Due to its significant LFC at both timepoints, the protein sequence for hypothetical protein m.24867 was interrogated using the protein Basic Local Alignment Search Tool (BLASTp) against the non-redundant protein database. The best-scoring descriptive annotation was "CyanoVirin-N domain containing protein" (CVNH: 95% Query coverage; e-value 2e-30; Fig. S4).

A second analysis of the mass spectrometry data was performed as a test to determine if known coral viral proteins could be detected from the whole-coral mass spectrometry analyses. Protein sequences from five viral proteomes that were translated from viral DNA that was isolated from corals (Table S2) (13). The resulting viral protein FASTA files were downloaded from Uniprot or NCBI (3.15.2022) and concatenated with the above mentioned *M. capitata* protein database, yielding a final database with 42,941 proteins (File S2). Mass spectrometry search parameters above were replicated with this database. A subset of the new ABACUS data with all viral proteins was generated with their respective spectral counts (148 unique viral proteins). Welches T-test was completed to determine if the number of viral proteins identified between $T_1R$ and $T_1S$ were significantly different. The same analysis was also completed on $T_2R$ and $T_2S$.

**MetaGOmics Biological Enrichment Analysis.** To determine if categories of proteins were enriched in the resilient vs. susceptible coral cohorts at the two timepoints, a biological enrichment strategy that analyzes Gene Ontology (GO) categorical terms was used to compare sets of detected proteins (14). Top results are reported with a cutoff E-value <1E-10. A fasta file of all *M. capitata* protein sequences confidently identified in these experiments (File S3) was analyzed with MetaGOmics v.0.1.1. MetaGOmics searched the truncated proteome against the UniProtKB Swiss-Prot non-redundant protein database (downloaded 10.15.2018) using BLASTp and assigned taxonomic and Gene Ontology (GO) terms for each protein sequence, and subsequently, to each peptide within the protein sequence. To determine biological enrichments of GO terms between experiments, peptides identified from each biological replicate from an experimental cohort were identified using percolator with an FDR cutoff <0.01 (15). Peptide spectral count values for each peptide that passed this threshold were averaged for the different experimental treatments ($T_1S$, $T_1R$, $T_2S$, $T_2R$). MetaGOmics generates laplace-corrected p-values to determine significance for log fold change calculations on the abundances of different GO terms. Here, we report Laplace corrected log fold change (base 2) results of all GO terms ≥0.5 and report a Laplace corrected Bonferroni corrected *p*-value significance <0.01 (Dataset S3A-B). MetaGOmics provides taxonomic-lowest common ancestor analyses and GO-based biological enrichment analysis of functions using Uniprot identifications. Although MetaGOmics was designed to analyze microbiomes, the use of the software was modified to work with a single organism by ignoring the taxonomic enrichment analysis to instead examine functions that are significantly enriched or depleted in pairwise comparisons of coral cohorts.

**Microbiome 16s rRNA Analyses.** Total DNA was extracted using the Qiagen DNA extraction kit (Qiagen). Half centimeter coral samples (N=12) were ground with a mortar and pestle in liquid nitrogen and then immediately transferred to 2 ml centrifuge tube containing buffer P1. Sample purity was measured using a NanoDrop ND1000 spectrophotometer (NanoDrop Technologies, Willmington DE). Samples showing good quality and concentrations were sent to Molecular Research LP (Shallowater, TX) for 16S rRNA amplicon sequencing. All 16S rRNA gene amplicon sequence data, processing steps and code for quality control on the microbiome data and analysis are available on GitHub (https://github.com/tanyabrown9/Resilient_vs_Susceptible_Mcapitata).

**16S rRNA Gene Amplicon Sequencing**. The V4 variable region of the 16S rRNA gene was amplified using the 515F (5´-GTGCCAGCMGCCGCGGTAA-3´) and 806R (5´-GGACTACHVGGGTWTCTAAT-3´) primer set (16). Amplification was conducted using a single-step 30-cycle PCR using HotStarTaq Plus Master Mix (Qiagen, USA) under the following conditions: 94°C for 30 seconds, 53°C for 40 seconds, and 72°C for 1 minute. A final elongation step at 72 °C was conducted for 1 minute. Sequencing was performed on an Ion Torrent PGM following manufacturers' guidelines at Molecular Research LP (Shallowater, TX). Sequences are deposited in NCBI as bioproject PRJNA933787. Initial sequencing resulted in a collection of 2,472,819 total reads, with an average read depth of 68,689 (± 28,107 SD) sequences per sample (Dataset S4A-B).

**Microbial Sequence Data Quality Analysis and Processing.** 16S rRNA gene amplicon sequence data from Molecular Research LP were processed using the QIIME2 software package (17). Forward and reverse reads from Molecular Research LP were imported into QIIME2 and demultiplexed (using the qiime paired-end sequencing method). Sequences were then denoised and chimaera-checked (using the qiime dada2 denoise-pyro method) to generate amplicon sequence variants (ASVs) (18). Care was taken to identify mitochondria and chloroplasts *in silico* prior to downstream analysis as previously described (19). Once annotated, chloroplast and mitochondria sequences were removed.

Alpha and beta diversity were assessed after the quality control steps. To conduct phylogenetic alpha and beta diversity metrics, a fragment insertion approach was used to insert short microbial 16S rRNA reads into a reference phylogeny. Sequences were placed on the QIIME2-compatible SILVA 128 tree included with the SEPP software using the QIIME fragment-insertion SEPP method (20, 21).

After rarefaction to 10,000 sequences per sample, phylogenetic-based alpha- and beta-diversity metrics were generated. Rarefaction was conducted to an equal depth to protect against false-positive results attributable to unequal sequencing depth between samples (22). Alpha diversity was assessed using the number of unique observed ASVs in rarefied samples (a measure of community richness) by the Simpson's Evenness and Shannon's Diversity Indexes. Overall differences in alpha diversity across susceptibility and time points were tested using Kruskal-Wallis tests. Post-hoc comparisons were performed within each group (*i.e.,* susceptibility and time point) as well as combined comparisons with *p*-values for pairwise tests between treatments adjusted for multiple comparisons using Bonferroni correction. The False Discovery Rate (FDR) for pairwise tests between susceptibility and time was controlled using FDR q-values reported by QIIME2. Beta diversity was assessed between samples using Weighted UniFrac distances and Bray-Curtis dissimilarities. The significance of differences in beta-diversity between susceptibility and time was tested using PERMANOVA (23).

All other statistical analyses of microbial taxonomy were conducted using R 4.0.3. The top 10 bacterial families in each sample type were selected for taxonomic analysis. Significant differences between bacterial families, susceptibility, and timepoint were carried out using a nested ANOVA. The Benjamini-Hochberg FDR was used to control the false discovery rate. Microbiome Multivariate Association with Linear Models was performed on the 16S data using the R package MaAsLin2 (24). Fixed effects for the analysis included bleaching status, resilience, time point, and whether the colony survived or succumbed to thermal stress (bleaching) by $T_2$. Coral colony term was set as a random effect and the analysis method was “LM”. Results were considered significant if the q-value was <0.05.

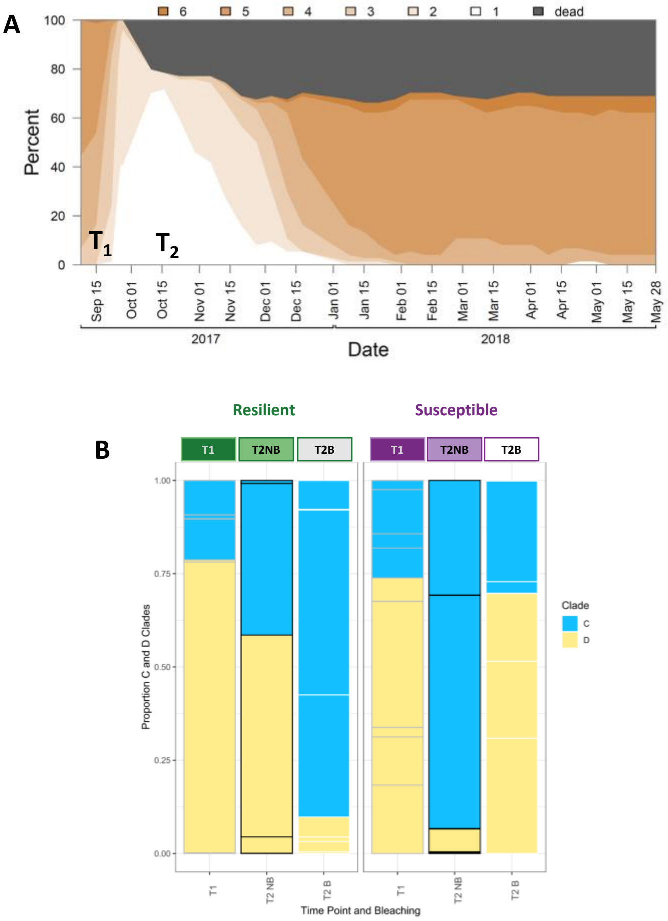


**Fig. S1**. A. Bleaching assessments were completed on all colonies every week using the Coral Watch Card where 1 indicates bleached, but not dead and 6 indicates the highest symbiont density. Noted times of T1 and T2 indicate when samples were collected for this study. B. Proportion of clade C (blue) and D (yellow) identified from Resilient (greens; *n=6*) and Susceptible (Purple; *n=6*) coral colonies at timepoint 1 (pre-bleaching) and timepoint 2 (NB- nonbleached cohort, B- thermally bleached cohort). No statistical differences were identified using ANOVA related to bleaching status, bleaching tolerance, or timepoint.

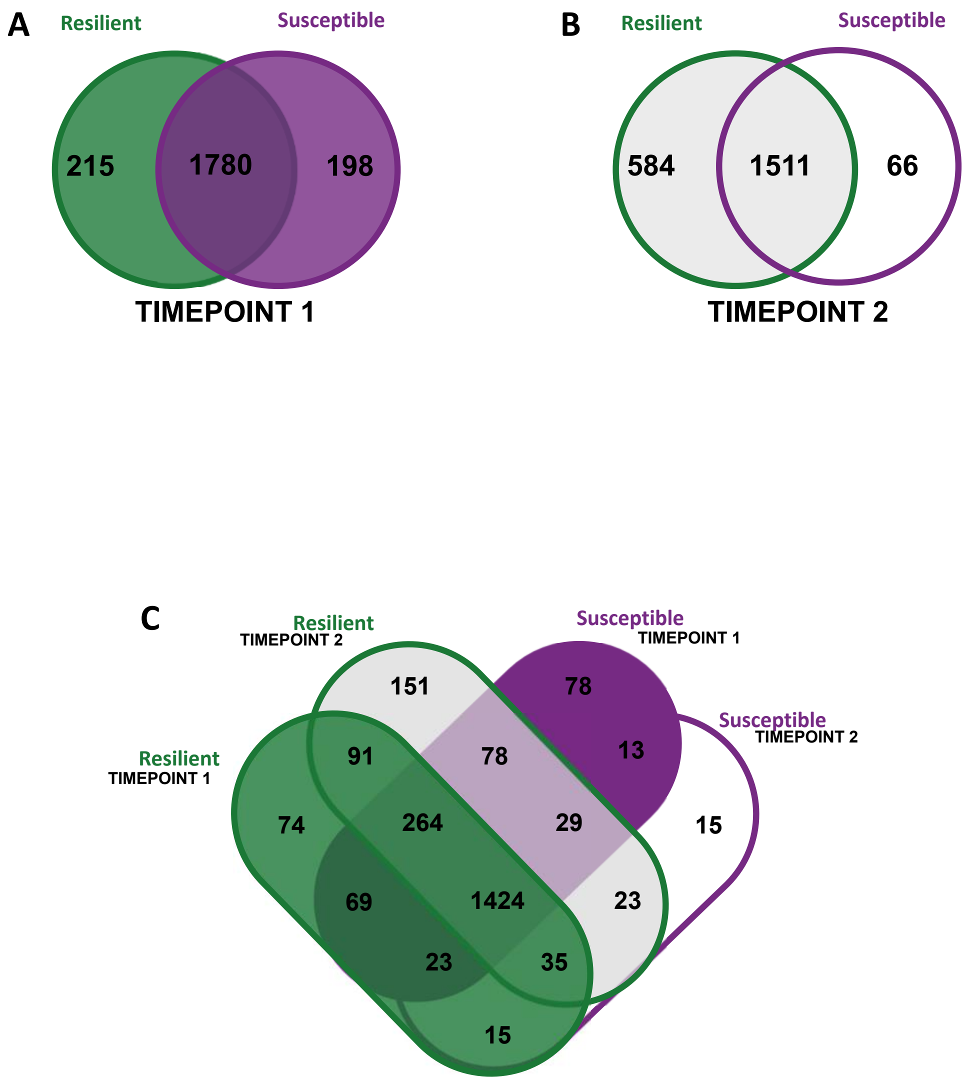


**Fig. S2.** Venn Diagram of proteins confidently Identified in resilient (*n=6*) and susceptible coral colonies (*n=6*) investigated at A. timepoint 1 before thermally bleached, B. timepoint 2 after thermal bleaching, and C. the overlap of all 4 treatments and timepoints.

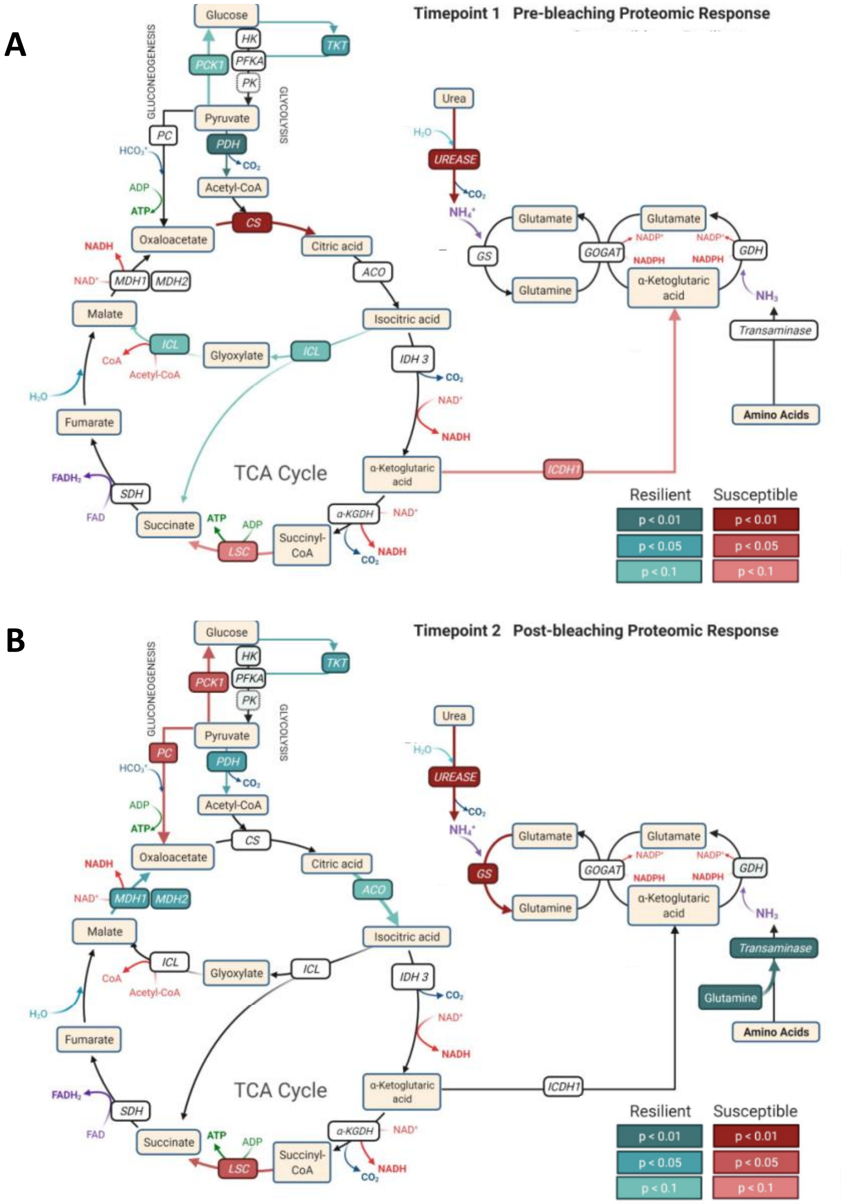


**Fig. S3.** Illustration of the proteins identified to be significantly increased in abundance in resilient (greens) or susceptible (reds) coral cohorts involved in the interconnected biochemical pathways of glycolysis/gluconeogenesis, the TCA cycle, urea degradation, and the glutamine synthase/glutamine oxoglutarate aminotransferase (GS/GOGAT) pathway at A. timepoint 1 before thermally bleached and B. timepoint 2 after thermal bleaching. Image was made using BioRender.

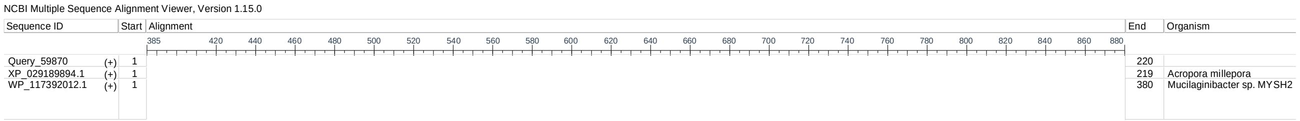


**Fig. S4.** Graphical illustration of a multiple sequence alignments of protein lcl|c238733_g2_i2|m.24867, noted to be significantly more abundant in the resilient colonies. The top query sequence in red (Query_59870) represents the input sequence mentioned. The next two sequences are from *Acropora millepora* and *Mucilaginbacter sp.* MYSH2 (both in red). Multiple alignment results revealed four highly conserved CyanoVirin-N domains (CVNH: 95% Query coverage; e-value 2e-30) depicted in grey.

**Table S1.** Resulting table from MaAsLin2 analysis. (24).

| TAXONOMY | FAMILY | METADATA | VALUE | COEF | STDERR | N | N.NOT.0 | PVAL | QVAL |
|---|---|---|---|---|---|---|---|---|---|
| **D_0__Bacteria.D_1__Proteobacteria.D_2__Gammaproteobacteria.D_3__Pseudomonadales.D_4__Moraxellaceae** | **Moraxellaceae** | **TimePoint** | **TimePoint** | -0.156 | 0.035 | 36 | 6 | 0.0001 | 0.005 |
| **D_0__Bacteria.D_1__Proteobacteria.D_2__Alphaproteobacteria.D_3__Caulobacterales.D_4__Caulobacteraceae** | **Caulobacteraceae** | **Resilience** | **S** | -0.411 | 0.089 | 36 | 6 | 0.0006 | 0.012 |
| **D_0__Bacteria.D_1__Proteobacteria.D_2__Gammaproteobacteria.D_3__Pseudomonadales. D_4__Moraxellaceae** | **Moraxellaceae** | **Resilience** | **S** | -0.233 | 0.062 | 36 | 6 | 0.0007 | 0.012 |
| **D_0__Bacteria.D_1__Proteobacteria.D_2__Alphaproteobacteria.D_3__Sphingomonadales.D_4__Sphingomonadaceae** | **Sphingomonadaceae** | **Resilience** | **S** | -0.181 | 0.062 | 36 | 7 | 0.0120 | 0.144 |
| **D_0__Bacteria.D_1__Firmicutes.D_2__Bacilli.D_3__Bacillales.D_4__Bacillaceae** | **Bacillaceae** | **Resilience** | **S** | -0.134 | 0.054 | 36 | 6 | 0.0194 | 0.186 |

**Table S2.** Table of coral viral proteomes selected, location the database was found, and the number of proteins downloaded.

| DESCRIPTION | DATABASE LOCATION | IDENTIFIER | # OF PROTEINS DOWNLOADED |
|---|---|---|---|
| *Cafeteria roenbergensis virus* | NCBI | NC_014637.1 | 1099 |
| *Paramecium bursaria Chlorella virus* | NCBI | NC_000852.5 | 1911 |
| *Lymphocystis disease virus* | NCBI | NC_005902.1 | 1488 |
| *Acanthamoeba polyphaga mimivirus* | UNIPROT | NC_014649.1 | 8727 |
| *Megavirus Chiliensis* | UNIPROT | NC_016072.1 | 4514 |

**Dataset 1. (separate file)** Physiological metrics on coral colonies from the study including: a readme file, symbiont density, symbiont clade distributions, chlorophyll a concentrations, lipid raw data, and lipid biomass data.

**Dataset 2. (separate file)** Processed proteomic data on each experiment in a range of formats: a readme file, proteins identified per experiment, Normalized Spectral Abundance Factors (NSAF) on all valid proteins identified, an accession number-based annotation file, QSPEC results for timepoints 1 and 2, ABACUS output file before processing.

**Dataset 3. (separate file)** MetaGOmics output files including: a readme file, MetaGOmics results from analysis of resistant vs. susceptible coral proteins identified at timepoint 1, MetaGOmics results from analysis of resistant vs. susceptible coral proteins identified at timepoint 2.

**Dataset 4. (separate file)** Microbiome sequence counts, metadata, and mapping information.

**File S1. (separate file)** Fasta files of protein sequences predicted from transcriptome for Montipora capitata plus contaminant protein database.

**File S2. (separate file)** Fasta files of protein sequences predicted from transcriptome for Montipora capitata plus contaminant protein database and 5 viral proteomes (see Table S1).

**File S3. (separate file)** Fasta file of all identified protein sequences from these experiments that were used as input for MetaGOmics analysis.
