## Supplemental figures for "Resilience in a time of stress: revealing the molecular underpinnings of coral survival following thermal bleaching events"

**This PDF file includes:**

Supporting text
Figures S1 to S4
Tables S1 to S2
Legends for Datasets S1 to S4
Legends for Files S1 to S3
SI References

**Other supporting materials for this manuscript include the following:**

Datasets S1 to S4
Files S1 to S3

Supplemental Methods

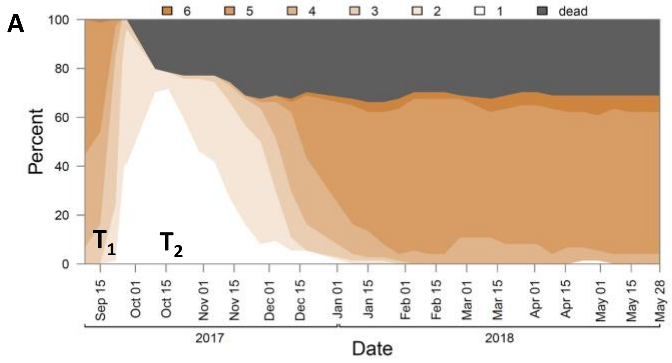


**Fig. S1**. A. Bleaching assessments were completed on all colonies every week using the Coral Watch Card where 1 indicates bleached, but not dead and 6 indicates the highest symbiont density. Noted times of T1 and T2 indicate when samples were collected for this study.

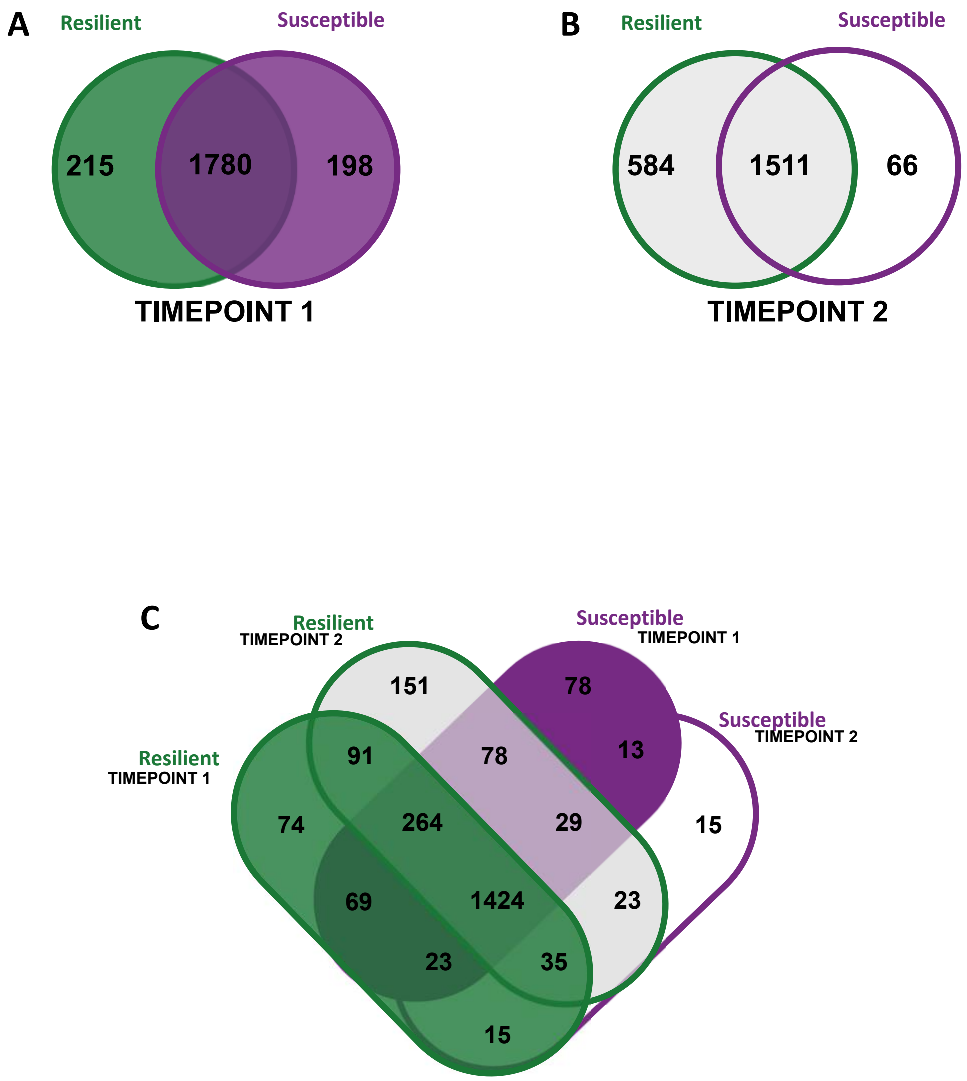


**Fig. S2.** Venn Diagram of proteins confidently Identified in resilient (*n*=6) and susceptible coral colonies (*n*=6) investigated at A. timepoint 1 before thermally bleached, B. timepoint 2 after thermal bleaching, and C. the overlap of all 4 treatments and timepoints.

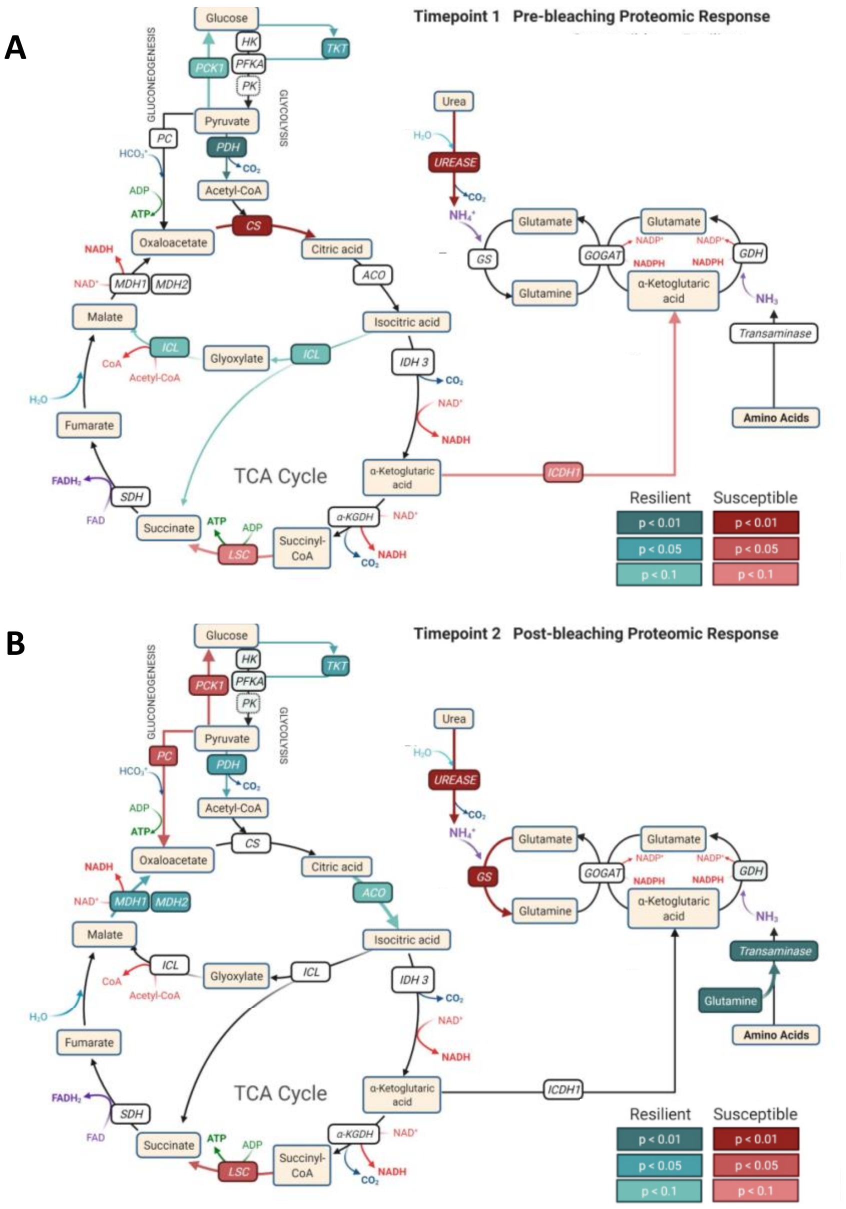


**Fig. S3.** Illustration of the proteins identified to be significantly increased in abundance in resilient (greens) or susceptible (reds) coral cohorts involved in the interconnected biochemical pathways of glycolysis/gluconeogenesis, the TCA cycle, urea degradation, and the glutamine synthase/glutamine oxoglutarate aminotransferase (GS/GOGAT) pathway at A. timepoint 1 before thermally bleached and B. timepoint 2 after thermal bleaching. Image was made using BioRender.

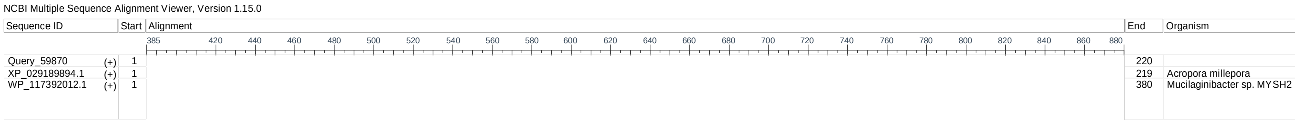


**Fig. S4.** Graphical illustration of a multiple sequence alignments of protein lcl|c238733_g2_i2|m.24867, noted to be significantly more abundant in the resilient colonies. The top query sequence in red (Query_59870) represents the input sequence mentioned. The next two sequences are from *Acropora millepora* and *Mucilaginbacter sp.* MYSH2 (both in red). Multiple alignment results revealed four highly conserved CyanoVirin-N domains (CVNH: 95% Query coverage; e-value 2e-30) depicted in grey.
